# Semaphorin3F reduces vascular endothelial and smooth muscle cell PI3K activation and decreases neointimal plaque formation

**DOI:** 10.1101/2022.03.22.485288

**Authors:** Chutima Rattanasopa, David Castano-Mayan, Chengxun Su, Aaron J. Farrugia, Maria Corlianò, Pakhwan Nilcham, Crystal Pang, Monalisa Hota, Koh Ser Mei, Wendy Lee, Dasan Mary Cibi, Atsu Aiba, Manvendra K. Singh, Siew Cheng Wong, Olaf Rotzschke, Alexander Bershadsky, Han Wei Hou, Elisa A. Liehn, Sujoy Ghosh, Roshni R. Singaraja

## Abstract

We previously conducted genetic analyses, and identified semaphorin signaling as associating with coronary artery disease. Of the semaphorins, human vascular expression profiling suggested *SEMA3F* as potentially linked to atherogenesis. In hyperlipidemic mice, SEMA3F reduced aortic lesion area, and increased fibrous cap endothelial content, leading to plaque stability. In a disturbed-flow-mediated endothelial dysfunction-driven lesion model, the absence of *Sema3f* increased plaques, further implicating SEMA3F in endothelial function. Monocyte adhesion to *Sema3f^-/-^* vascular endothelial cells (VECs) was elevated, driven by increased PI3K activity, leading to increased NF-κB-mediated elevation in VCAM1 and ICAM1 expression, suggesting that SEMA3F reduces VEC PI3K activity. Increased permeability led to increased monocyte transmigration through *Sema3f^-/-^* VECs, and mTOR phosphorylation was decreased, suppressing VE-cadherin expression and cell-cell adherens junction stability. Actomyosin fiber formation was decreased in *Sema3f^-/-^* VECs, which was reversed by PI3K inhibition, further implicating SEMA3F in adherens junction stability. In *Sema3f^-/-^* vascular smooth muscle cells (VSMCs), active PI3K was also increased. PI3K facilitates VSMC proliferation, migration, and pro-atherogenic phenotype switching, which were reduced by SEMA3F. In agreement, in a model of VSMC proliferation and migration-induced neointima formation, SEMA3F reduced plaques. Semaphorin3F is causally atheroprotective. SEMA3F’s suppression of VEC and VSMC PI3K activation may contribute to its atheroprotection.

## Introduction

Atherosclerotic coronary artery disease (CAD) is the primary cause of ischemic heart disease, and a leading cause of death worldwide (1). Currently available treatment strategies aim to reduce risk factors via lifestyle changes, decrease low density lipoprotein cholesterol (LDL-C) via statins and proprotein convertase subtilisin/kexin type 9 (PCSK-9) inhibitors, blood pressure surveillance, antithrombotic and anti-inflammatory drugs, and surgical interventions (2, 3). However, mortality due to CAD remains high (1). Therefore, additional mechanism-based therapies are urgently needed to target atherosclerotic CAD more directly.

Previously, we used an integrative systems genetics approach on a large scale genome-wide association meta-analysis (>66,000 controls, >24,000 CAD cases from the CARDIoGRAM consortium) and identified 32 biological pathways as being significantly associated with CAD (4). Notably, the integrative analysis revealed semaphorin signaling as one of the most significant CAD-associated pathways (p<0.001). Semaphorins are a family of secreted or membrane-associated effectors, with a signaling network composed of ligands (Semaphorins), receptors (Plexins), co-receptors (Neurophilins), intracellular effectors (Crmps/DPYSLs), and regulator kinases (FES, FYN, CDK5, GSK3B). Widely studied in neuronal development and cancers, the involvement of semaphorin signaling in heart development is also well established (5).

The literature around the direct role of semaphorins in atherosclerosis is more modest, being largely correlative in nature and primarily based on in vitro observations that do not allow for causal inference. Our study builds on, but is distinct from these earlier reports, and expands the clinical relevance of semaphorins in several important ways. Through pathway-based inference of genetic association data, we demonstrated an association between semaphorin gene polymorphisms in humans and CAD (4). Through *in-vivo* knockout studies and studies administering SEMA3F, here we establish a causal relationship between SEMA3F and vascular disease and interrogate the underlying mechanisms. By employing multiple models of vascular dysfunction, our findings expand the clinical scope of SEMA3F as a possible therapeutic target for atherosclerosis (Supplemental Figure 1). Together, these findings create a fuller picture of the complexity of semaphorin biology in vascular dysfunction, and advance SEMA3F as a candidate therapeutic target for atheroprotection.

## Results

### *SEMA3F* is expressed in human atherosclerosis-relevant tissues

We identified ‘Crmps in semaphorin signaling’ in a large scale genome-wide association meta-analysis as being significantly associated with CAD (4). To determine which member(s) of the semaphorin family were expressed in atherosclerosis relevant tissues in humans, we compiled expression data from the publicly available Gene Expression Omnibus (GEO) (6). *SEMA3F* expression was substantially higher compared to most other semaphorins in vascular endothelial (VECs) and *SEMA3F* also showed high expression in smooth muscle cells (VSMCs) and monocytes/macrophages (Supplemental Figure 2, A-C), all cell types with critical roles in atherosclerosis, as well as in human atherosclerotic plaques (Supplemental Figure 2D). Of the class 3 semaphorins, *SEMA3F* was also the most highly expressed in endothelial cell lines (7). Although *SEMA3C* was the only other family member highly expressed in vascular tissues, *Sema3c^-/-^* mice die within a few hours after birth (8), making the study of the role of SEMA3C in CAD difficult. To determine if SEMA3F was expressed in the vasculature of mice, immunohistological staining of aortas in chow-fed *Ldlr^-/-^* mice was performed. SEMA3F was robustly expressed in endothelial cells (VE-cadherin^+^), smooth muscle cells (SMA^+^), and macrophages (MAC2^+^) (Supplemental Figure 2E). These data suggest that *SEMA3F* is highly expressed in vascular tissue and may modulate coronary artery disease.

### SEMA3F administration reduces arterial plaques

We next determined if SEMA3F modulates arterial lesion formation by administering SEMA3F protein to western-type diet (WTD)-fed *Ldlr^-/-^* mice. Aortic sinus lesion area was decreased by ∼30% in SEMA3F-administered mice (Control: 12.1±1.3; SEMA3F: 8.1±0.8, % total lesion area/total aortic sinus area, n=9 each, p=0.02) (Figure 1A). This was driven by a ∼65% reduction in advanced lesion area (Control: 45.8±11.0; SEMA3F: 16.5±7.3, % lesion area/total lesion area, n=9 each, p=0.04), as categorized by the American Heart Association classification system (9), whereas mild lesion area was increased (Control: 54.2±11.0; SEMA3F: 83.5±7.3, % lesion area/total lesion area, n=9 each, p=0.04) (Supplemental Figure 3A) suggesting that SEMA3F decreased the progression of mild to more severe lesions. Neutral lipid-positive area in *en face* stained aortic arches were also decreased in SEMA3F-treated mice (Control: 17.1±1.5; SEMA3F: 12.0±0.9, % lesion area/aortic area, n=9 each, p=0.01) (Supplemental Figure 3, B and C). SEMA3F did not alter body weights or plasma lipoprotein levels (Supplemental Figure 4, A-D). These data show that SEMA3F is atheroprotective, and reduces the progression of mild to advanced lesions in a background of hyperlipidemia.

**Figure 1:**
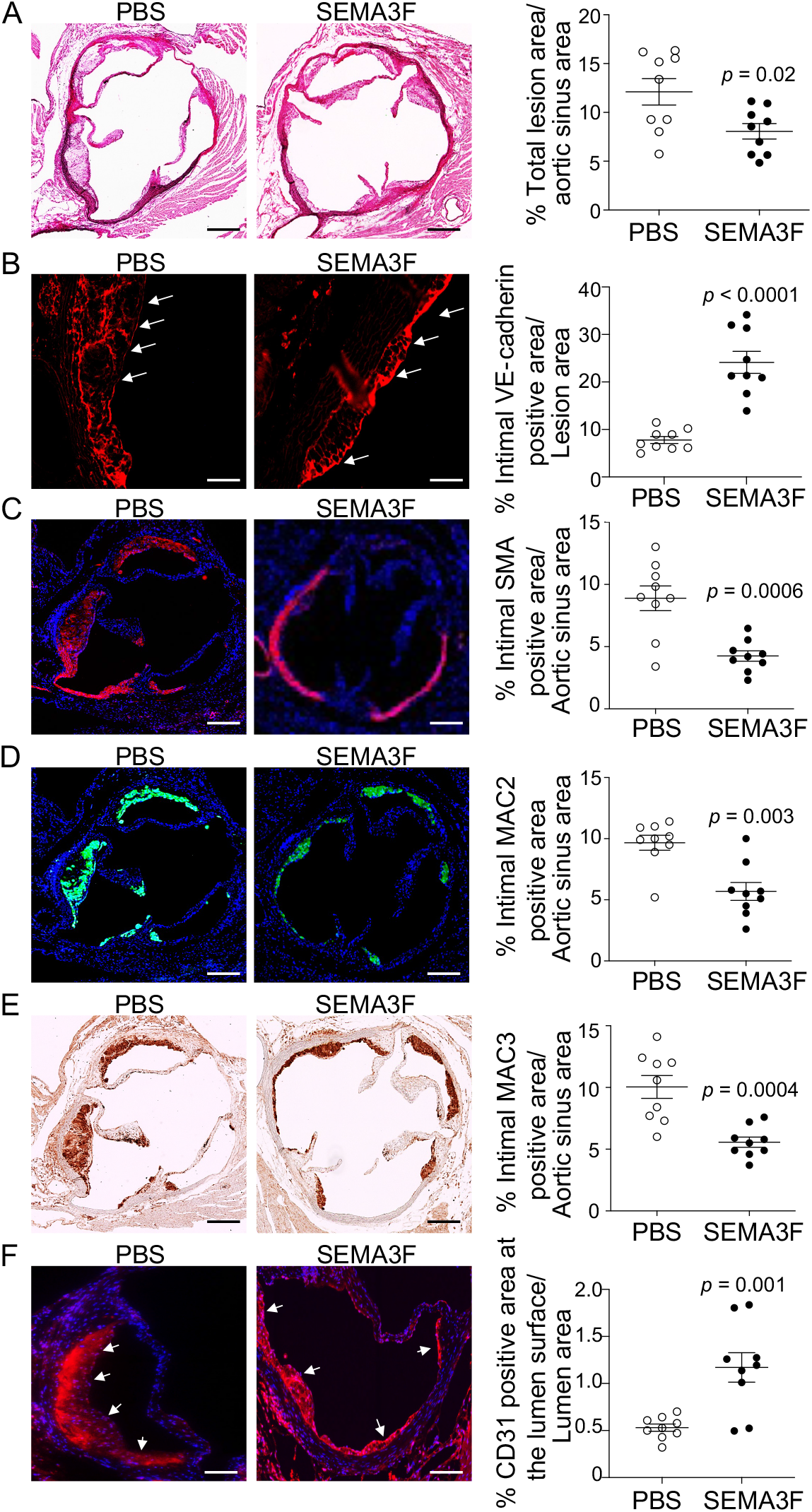
SEMA3F administration reduces atherosclerotic lesion area and increases lesion stability in hyperlipidemic mice. **(A)** Representative photomicrographs of hematoxylin-phloxine-saffron stained aortic sinus showing that total atherosclerotic lesion area was decreased in SEMA3F-treated western-type diet-fed *Ldlr^-/-^* mice. Increased **(B)** VE-cadherin positive area (red) in the luminal surface of lesions, and decreased **(C)** α-smooth muscle actin (SMA) positive area (red), **(D)** MAC2 positive area (green), and **(E)** MAC3 positive area (brown) in SEMA3F-treated mice. DAPI (blue). **(F)** Increased CD31 positive endothelial area (red) in the luminal surface of lesions. Groups abbreviated as: PBS, *Ldlr^-/-^* mice fed western-type diet, injected with PBS; SEMA3F, *Ldlr^-/-^* mice fed western-type diet, injected with SEMA3F protein. Values are mean±SEM, 6 sections/mouse, n=9/group. Data in (A) to (C), (E) and (F) were normally distributed and analyzed using Student’s *t*-tests. Data in (D) were not normally distributed and analyzed using Mann Whitney *U*-test. Scale bar in (A, C-E): 200 µm, (B) 50 µm, (F) 100 µm.

### SEMA3F administration increases plaque stability

We next assessed lesional endothelial, smooth muscle and macrophage content to determine how SEMA3F might reduce lesion formation. VE-Cadherin staining showed that SEMA3F administration increased intimal endothelial area by ∼300% (Figure 1B) (Control: 7.8±0.7; SEMA3F: 24.1±2.3, n=9 each; % intimal VE-cadherin positive area/lesion area, p<0.0001), suggesting increased plaque stability. α-smooth muscle actin (SMA) (10) immunostaining showed ∼50% decrease in lesion smooth muscle area (Control: 8.9±1.0; SEMA3F: 4.3±0.4, n=9 each; % intimal SMA positive area/aortic sinus area, p=0.0006) (Figure 1C), and both activated (MAC2) (Control: 9.7±0.6; SEMA3F: 5.7±0.7, n=9 each, % intimal MAC2 positive area/aortic sinus area, p=0.003) (Figure 1D), and non-activated (MAC3) (Control: 10.0±0.9; SEMA3F: 5.6±0.4 n=9 each, % intimal MAC3 positive area/aortic sinus area, p=0.0004) (Figure 1E) macrophage positive areas were decreased in SEMA3F-treated mice. The increase in endothelial area, assessed by staining for VE-cadherin, a marker specific for vascular endothelial cells (11) was confirmed by staining for CD31, a marker for VECs that can cross react with leukocytes (12), and found to be increased in the luminal surface of the lesions (Control: 0.53±0.04; SEMA3F: 1.17±0.16, n=9 each, % intimal MAC2 positive area/aortic sinus area, p=0.001) (Figure 1F). These data suggest that SEMA3F administration, in this model, decreases plaque cellularity and macrophage content, while increasing plaque stability, as evidenced by the increased VE-cadherin positive endothelial layer in the luminal surface of the plaque.

### Increased lesion area in *Sema3f^-/-^* mice with carotid partial ligation

SEMA3F regulates endothelial cell function, and modulates monocyte migration in umbilical vein cells (7). We found increased endothelial staining in the luminal surface of plaques in SEMA3F-administered mice. Together, these data suggest that SEMA3F may modulate VEC function to affect lesion formation. Partial ligation of the blood vessels increases turbulent flow, and models endothelial cell activation response to disturbed flow, which, on a background of hypercholesterolemia, rapidly increases atherogenesis in mice (13). We quantified lesion area in WTD-fed *Sema3f^-/-^* mice that had undergone partial ligation of their common carotid artery (13). Carotid lesion area was increased in *Sema3f^-/-^* mice (*WT*: 22.8±2.8; *Sema3f^-/-^*: 39.2±3.9; % intimal area/total vessel area, n=8 each, p=0.004) (Figure 2A). While body weights of *Sema3f^-/-^* mice were decreased (*WT*: 32.7±1.1; *Sema3f^-/-^*: 21.7± 0.4; gram, n=8 each, p<0.0001) (Supplemental Figure 5A), lesion measures were normalized to total vessel area to account for any size differences between mice. Intimal endothelial cell area was decreased by ∼30% in *Sema3f^-/-^* mice (*WT*: 63.2±4.6; *Sema3f^-/-^*: 45.7±4.4, n=8 each; % intimal VE-cadherin positive area/vessel area, p=0.01) (Figure 2B), in line with our finding that SEMA3F administration increased VE-cadherin positive endothelial area. The number of smooth muscle cells (α-SMA^+^) in the intimal lesions was increased ∼4-fold in *Sema3f^-/-^* mice (*WT*: 29.0±7.6; *Sema3f^-/-^*: 121.0±12.5; n=8 each, p=0.0003) (Figure 2C). Lesional macrophage (MAC2^+^) numbers were unchanged in this model at this time-point (*WT*: 0.5±0.3; *Sema3f^-/-^*: 0.6±0.3; n=8 each, p=0.9) (Figure 2D). These mice are not on the *Ldlr^-/-^* background, and therefore not hyperlipidemic, suggesting that hyperlipidemia-induced macrophage activation (14) may not contribute to lesions in these mice. As well, in this model, lesion development was assessed at only 5 weeks after carotid partial ligation. Assessing lesional macrophage content at later time-points may result in altered lesional macrophage numbers. Together, these data show that the absence of SEMA3F increases intimal plaque area and cellularity in a mouse model of disturbed flow-mediated endothelial cell activation, whereas endothelial positive area was decreased, suggesting that endothelial cell SEMA3F contributes to lesion development. Plasma total cholesterol levels were decreased (Supplemental Figure 5B) in *Sema3f^-/-^* mice, driven by a decrease in HDL cholesterol levels (Supplemental Figure 5C), whereas plasma non-HDL cholesterol and triglyceride levels were unchanged (Supplemental Figure 5, D and E). However, SEMA3F is unlikely to affect lesions via altering plasma HDL cholesterol levels for two reasons. First, SEMA3F did not affect plasma HDL-C levels in the hyperlipidemic SEMA3F administered mice (Supplemental Figure 4C), although lesion area was altered in these mice. Second, flux through the HDL pathway (HDL functionality), rather than the absolute circulatory HDL-cholesterol levels maybe more robustly associated with atherosclerotic cardiovascular disease (ASCVD), and a causative association between plasma HDL-cholesterol levels and ASCVD has not been established (15).

**Figure 2:**
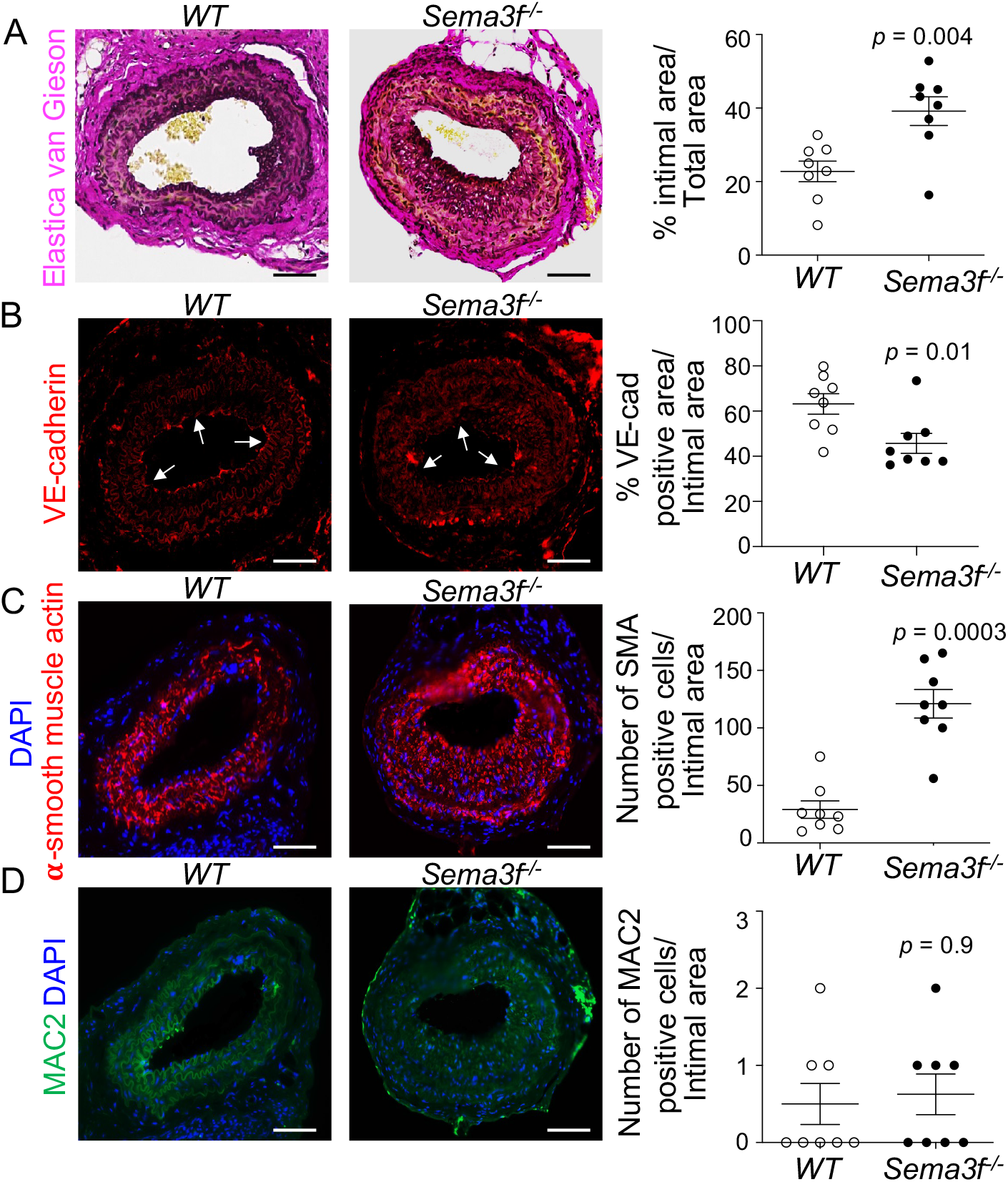
Increased arterial lesion area in *Sema3f^-/-^* mice. **(A)** Increased lesion area in elastica van Gieson-stained carotid sections in *Sema3f^-/-^* mice with partially ligated common carotid artery. **(B)** Decreased VE-cadherin positive endothelial cell area (red), **(C)** increased smooth muscle actin positive (SMA, red) vascular smooth muscle cells, and **(D)** unchanged MAC2 positive (green) macrophages in *Sema3f^-/-^* mice. DAPI (blue). Groups abbreviated as: *WT*, partially ligated *wild-type* mice fed western-type diet; *Sema3f^-/-^*, partially ligated *Sema3f* knockout mice fed western-type diet. Values are mean±SEM, 6 sections/mouse, n=8/group. Data in (A) were normally distributed and analyzed using Student’s *t*-test. Data in (B) to (D) were analyzed using Mann-Whitney *U*-tests. Scale bar: 50 µm.

### Vascular endothelial cells from *Sema3f^-/-^* mice show increased monocyte adhesion

To determine if SEMA3F expression is altered during VEC activation, we treated primary human coronary artery endothelial cells (HCAECs) with Tumour Necrosis Factor alpha (TNF-*α*) (16), and found reduced SEMA3F expression (Control: 1.08±0.03; TNF-*α*: 0.73±0.04, relative fold change/β-actin, n=3 each, p=0.002) (Supplemental Figure 6A). To confirm that the HCAECs were indeed activated by TNF-*α*, we assessed Vascular cell adhesion molecule-1 (VCAM1) and Intercellular adhesion molecule-1 (ICAM1) levels post TNF-*α* treatment, and found increased VCAM1 (Control: 0.86±0.003; TNF-*α*: 1.06±0.01, relative fold change/β-actin, n=3 each, p<0.0001) and ICAM1 expression (Control: 1.07±0.02; TNF-*α*: 1.25±0.01, relative fold change/β-actin, n=3 each, p=0.001) (Supplemental Figure 6A). These data suggest that endothelial SEMA3F may be atheroprotective. Thus, we assessed the monocyte adhesion capability of VECs. Both unstimulated THP-1 (Supplemental Figure 6B, ANOVA p=0.001) and mouse splenic monocyte (Figure 3A, ANOVA p=0.002) adhesion to *Sema3f^-/-^* VECs were increased by 60%. These increases in THP-1 and mouse monocyte adhesion were fully reversed upon pre-treatment of *Sema3f^-/-^* VECs with purified SEMA3F protein, which was washed off before the flow of monocytes was initiated (Figure 3A, ANOVA p=0.002 and Supplemental Figure 6B, ANOVA p=0.001), suggesting that increased monocyte adhesion in the absence of SEMA3F in this model resulted solely from the modulation of VEC function by SEMA3F.

**Figure 3:**
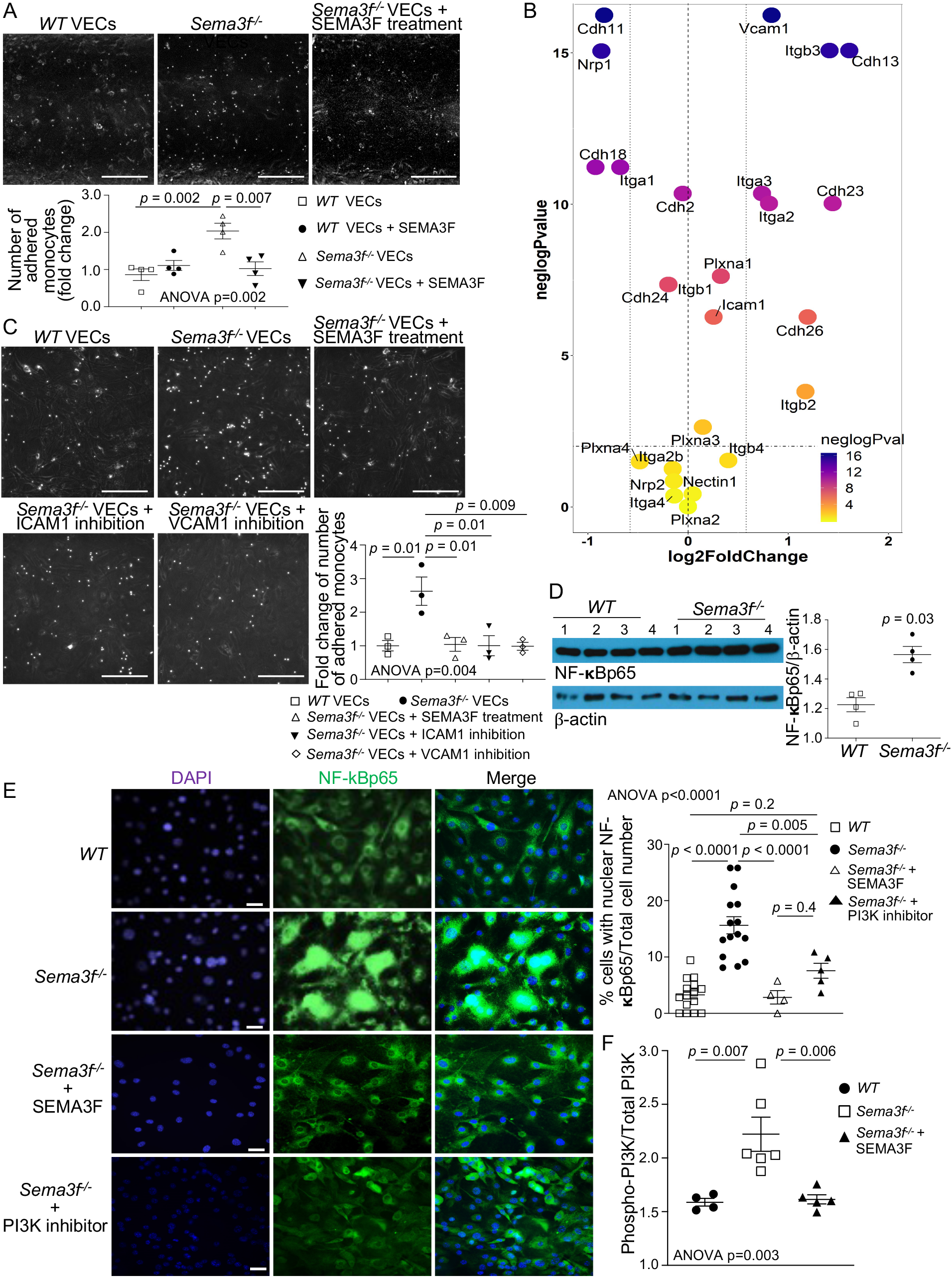
PI3K/NF-κB-mediated increased endothelial cell adhesion molecule expression underlies increased monocyte adhesion to *Sema3f^-/-^* endothelial cells. **(A)** The number of human monocytes adhered to *Sema3f^-/-^* VECs was higher, and pre-treating *Sema3f^-/-^* VECs with SEMA3F protein reversed monocyte adhesion, showing that SEMA3F directly modulates endothelial monocyte adhesion. n=4/group. **(B)** RNA sequencing identified increased *Vcam1* and *Icam1* adhesion molecule expression in *Sema3f^-/-^* aortic VECs. **(C)** Increased adhesion of mouse splenocytes to *Sema3f^-/-^* VECs, which was reduced upon antibody blocking of ICAM1 or VCAM1, showing that increased VCAM1/ICAM1 expression in *Sema3f^-/-^* VECs drove the increased monocyte adhesion. n=3/group. **(D)** Increased activation of the VCAM1/ICAM1 regulator NF-κB, assessed by NF-κBp65 levels in *Sema3f^-/-^* aorta. **(E)** The p65 fragment of NF-κB (green), which represents its activated state, shows significantly increased nuclear localization in isolated aortic endothelial cells from *Sema3f^-/-^* mice, which was fully reversed upon treatment of the *Sema3f^-/-^* cells with SEMA3F protein, as well as with the PI3K inhibitor Wortmannin. DAPI = blue. **(F)** Increased levels of active PI3K (phospho-tyrosine 467/199 PI3K p85) in aortic endothelial cells from *Sema3f^-/-^* mice, which was fully reversed upon treatment of the *Sema3f^-/-^* cells with SEMA3F protein. Groups are abbreviated as: *WT*, wild type; *Sema3f^-/-^*, *Sema3f* knockout mice; VECs, vascular endothelial cells. Values represent mean±SEM. Data in (A), (C), (E) and (F) were analyzed using One-way ANOVA followed by Tukey post-hoc test for multiple comparisons. Data in (B) were analyzed in R using *limma*. Data in (D) were assessed using Mann-Whitney *U*-tests. Scale bar in (A) and (C): 200 µm, (E): 100 µm.

### Vascular endothelial cells from *Sema3f^-/-^* mice show increased cell surface adhesion molecule expression

Increased monocyte adhesion to VECs maybe modulated by increased adhesion molecule expression in either activated monocytes or in VECs, or both. However, in the above experiments, the adhesion of unstimulated monocytes was assessed, and pre-treatment of *Sema3f^-/-^* VECs with SEMA3F protein reversed the increased monocyte adhesion, suggesting that endothelial SEMA3F modulated the monocyte binding.

To determine if SEMA3F modulated global changes in VEC adhesion molecule expression, we performed RNA sequencing of *Sema3f^-/-^* VECs and found 598 genes differentially expressed at absolute log fold-change *≥*1 and adjusted p-value *≤*0.05. 226 genes were down-regulated and 372 genes were up-regulated, of which the top 30 are shown (Supplemental Figure 7). The KEGG pathway relating to cell-adhesion genes (‘Cell adhesion molecules’) was up-regulated (p_adj_=0.001), whereas the ‘Extracellular matrix (ECM)-receptor interaction’ pathway involving multiple collagen genes was down-regulated (p=0.005) (Supplemental Figure 8). Within this pathway, the expression of adhesion molecules *Vcam1* and *Icam1,* which mediate immune cell rolling, adhesion and extravasation (17), was upregulated in *Sema3f^-/-^* VECs (p<0.0001) (Figure 3B). Western immunoblots confirmed the increased VCAM1 (*WT*: 0.95±0.02; *Sema3f^-/-^*: 1.42±0.03, fold change, normalized to β-actin, n=4 each, p=0.03) and ICAM1 (*WT*: 1.37±0.02; *Sema3f^-/-^*: 1.81±0.03, fold change, normalized to β-actin, n=4 each, p=0.03) (Supplemental Figure 9) in *Sema3f^-/-^* aortae.

To determine if increased VCAM1 and ICAM1 in *Sema3f^-/-^* VECs facilitated the increased monocyte adhesion, *Sema3f^-/-^* VECs were treated with anti-VCAM1 or anti-ICAM1 antibodies, which resulted in reduced monocyte adhesion to *Sema3f^-/-^* VECs (*WT*: 1.00±0.15; *Sema3f^-/-^*: 2.63±0.42; *Sema3f^-/-^* +ICAM1 antibody: 1.00±0.30; *Sema3f^-/-^* +VCAM1 antibody: 0.99±0.12, fold change, n=3 each, ANOVA p=0.004) (Figure 3C). Thus, the increased monocyte adhesion to *Sema3f^-/-^* VECs was modulated by increased cell-surface ICAM1 and VCAM1 expression.

Both VCAM1 and ICAM1 are induced by the activation of nuclear factor kappa-light-chain-enhancer of activated B cells (NF-κB) and the p65 sub-unit of NF-κB is increased upon its activation (18). NF-κB p65 levels were increased in *Sema3f^-/-^* aorta (*WT*: 1.23±0.05; *Sema3f^-/-^*: 1.57±0.06, fold change, normalized to β-actin, n=4 each, p=0.03) (Figure 3D). Active NF-κB p65 increases *VCAM1* and *ICAM1* transcription (18). Compared to its mostly cytoplasmic localization in *wild-type* aortic VECs, NF-κB p65 in *Sema3f^-/-^* aortic VECs showed increased nuclear localization (*WT*: 3.3±0.7; *Sema3f^-/-^*: 15.6±1.5, % cells with nuclear NF-κB/total cell number per field, n=15 each, ANOVA p<0.0001), which was reversed by pre-treatment of *Sema3f^-/-^* VECs with SEMA3F protein (*Sema3f^-/-^*: 15.6±1.5, n=15; *Sema3f^-/-^* + SEMA3F: 2.8±1.2, n=4, % cells with nuclear NF-κB/total cell number per field, ANOVA p<0.0001) (Figure 3E). Thus, increased activated NF-κB p65 in *Sema3f^-/-^* VECs increases VCAM1 and ICAM1 levels, leading to increased monocyte adhesion to the VECs.

NF-κB is activated by the Phosphoinositide 3-kinase (PI3K) pathway (19), and SEMA3F has been suggested to inhibit PI3K in venous cells (20). To determine if the increased NF-κB activation in *Sema3f^-/-^* VECs was modulated by increased PI3K, we assessed NF-κB p65 localization in response to PI3K inhibition, and found that the increased nuclear localization of NF-κB p65 in *Sema3f^-/-^* VECs was reversed upon treatment with Wortmannin, an inhibitor of PI3K (21) (*Sema3f^-/-^*: 15.6±1.5, n=15; *Sema3f^-/-^* + PI3K inhibitor: 7.6±1.3, n=5, % cells with nuclear NF-κB/total cell number per field, ANOVA p<0.0001) (Figure 3E).

These findings suggest that in *Sema3f^-/-^* VECs, increased PI3K activity drives the increase in NF-κB p65 activation and its nuclear localization, thus increasing VCAM1/ICAM1 expression. Thus we assessed PI3K activity in *Sema3f^-/-^* VECs, by quantifying phospho-tyrosine 467/199 PI3K p85 levels (22) and found it increased (*WT*: 1.59±0.04, n=4; *Sema3f^-/-^*: 2.22±0.16, n=6; Phospho-PI3K/Total PI3K ratio, ANOVA p=0.003) (Figure 3F). Treatment of *Sema3f^-/-^* VECs with SEMA3F reversed this increased PI3K activation (*Sema3f^-/-^*: 2.22±0.16, n=6; *Sema3f^-/-^* + SEMA3F: 1.62±0.04, n=5; Phospho-PI3K/Total PI3K ratio, ANOVA p=0.003) (Figure 3F), indicating that the increased PI3K activity resulted from the absence of SEMA3F. Our findings suggest that the PI3K/NF-κB pathway contributes to increased monocyte adhesion to VECs in the absence of SEMA3F.

### Monocyte transmigration through *Sema3f^-/-^* vascular endothelial cells is increased

We next assessed monocyte transmigration through *Sema3f^-/-^* VECs. THP-1 monocyte migration through *Sema3f^-/-^* VECs was increased (*WT*: 4.4±0.9; *Sema3f^-/-^*: 8.9±0.9 x10^5^ cells/mL, n=9 each, p=0.006) (Supplemental Figure 10). Since THP-1 monocytes are human derived, we performed this experiment using splenic cells isolated from *wild-type* mice, and again found increased transmigration through *Sema3f^-/-^* VECs (*WT*: 1.5±0.2; *Sema3f^-/-^*: 2.6±0.2, x10^6^ cells/mL, n=6 each, p=0.006) (Figure 4A), suggesting that the increased transmigration may have resulted from increased leakage of the *Sema3f^-/-^* endothelial cell layer.

**Figure 4:**
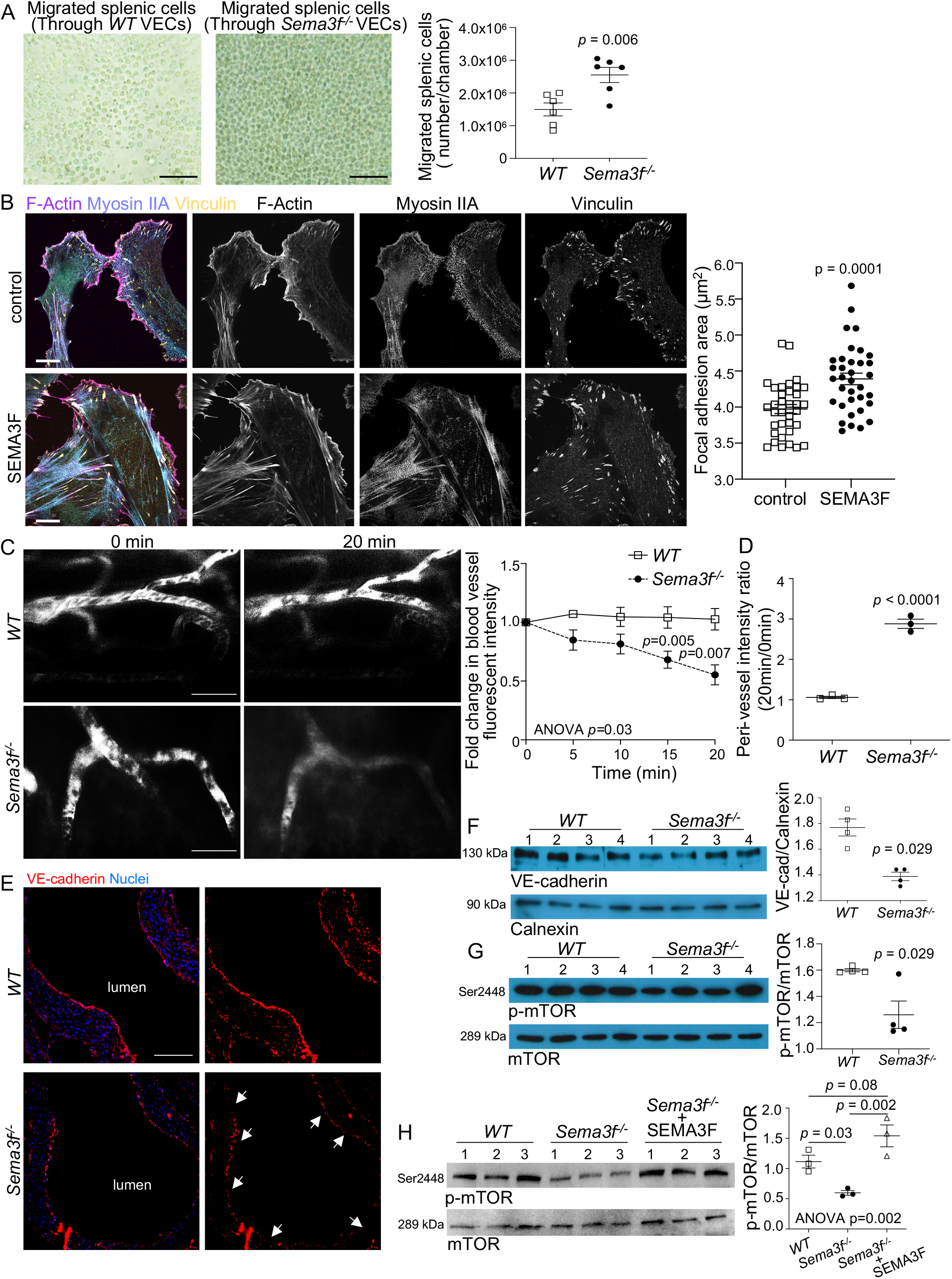
SEMA3F modulates endothelial cell F-actin dynamics, VE-cadherin expression, and vascular endothelial cell membrane barrier integrity. **(A)** The number of *WT* mouse splenic cells migrated through *Sema3f^-/-^* VECs is higher than through *WT* VECs. n=6/group. **(B)** Increased actomyosin fibers (purple and blue) and focal adhesion area (yellow) in primary human aortic endothelial cells treated with SEMA3F. n=35/group. **(C)** Reduced FITC-dextran fluorescence intensity in ear blood vessels and **(D)** increased FITC-dextran in the peri-vascular region of *Sema3f^-/-^* mice, assessed by intravital two-photon microscopy. n=3/group. **(E)** Representative photomicrographs of VE-cadherin (Red) and DAPI (Blue) showing disorganized endothelial barrier in *Sema3f^-/-^* aorta. **(F)** Reduced VE-cadherin in *Sema3f^-/-^* aorta, n=4/group. **(G)** Phospho-mTOR levels are decreased in the aorta of *Sema3f^-/-^* mice. n=4/group. **(H)** Phospho-mTOR is also decreased in isolated aortic VECs from *Sema3f^-/-^* mice, which is reversed upon treatment of *Sema3f^-/-^* VECs with SEMA3F. n=3/group. Groups abbreviated as: *WT*, wild type mice; *Sema3f^-/-^*, *Sema3f* knockout mice. VECs, vascular endothelial cells. Values are mean±SEM. Data in (A), (B) and (D) were analyzed using Student’s *t*-tests. Data in (C) were analyzed using Two-way ANOVA. Data in (F) and (G) were assessed using the Mann-Whitney *U*-tests. Data in (H) were analyzed using One-way ANOVA followed by Tukey post-hoc tests for multiple comparisons. Scale bar in (A), (C): 100 µm, (B): 10 µm, (D): 25 µm.

### *Sema3f^-/-^* mice show vascular endothelial remodeling and reduced endothelial barrier integrity

F-actin modulates cell-cell adherens junction dynamics, and VE-cadherin regulates endothelial cell adherens junction stability, thus modulating vascular permeability (23). SEMA3F was shown to modulate endothelial permeability in human umbilical vein cells via regulating F-actin filaments (7). However, umbilical vein cells are not an ideal model to assess atherosclerotic mechanisms since they do not undergo high shear stress, nor do they show similar cell junctions compared to aortic endothelial cells (24). Thus, we assessed actomyosin filaments and adherens junction dynamics in SEMA3F-treated primary human coronary artery endothelial cells (HCAECs), a more physiological model for studying atherosclerosis. We found increased actomyosin stress fibers and larger focal adhesions upon SEMA3F administration, hallmarks of RHO activation (Control: 3.97±0.06; SEMA3F: 4.39±0.08, focal adhesion area (μm^2^), n=35 each, p=0.0001, Figure 4B). While severe activation of RHO accompanied by strong augmentation of stress fibers and focal adhesions may result in the rupture of cell-cell contacts and increased endothelial permeability (25), it is well documented that contractility-driven moderate tugging force can in fact reinforce cell-cell junctions and promote endothelial barrier function (26). Therefore, we assessed the integrity of VE-cadherin junctions in confluent HCAECs treated with SEMA3F. In line with the moderate increase in actomyosin filaments observed, SEMA3F treatment promoted thicker VE-cadherin junctions in cell monolayers (Control: 1.43±0.05; SEMA3F: 1.93±0.06, VE-cadherin junction width (μm), n=111 each, p<0.0001, Supplemental Figure 11), again suggesting that SEMA3F decreases endothelial permeability.

PI3K signaling decreases actomyosin contractility in endothelial cells (27). To confirm if SEMA3F-mediated decrease in PI3K activation contributed to the increased F-actin stress fibers, we performed F-actin staining in *Sema3f^-/-^* VECs treated with the PI3K inhibitor Wortmannin. F-actin fibers were decreased in *Sema3f^-/-^* aortic VECs, which was reversed upon treatment of the *Sema3f^-/-^* VECs with Wortmannin (Supplemental Figure 12), suggesting that the increased PI3K activation in *Sema3f^-/-^* VECs resulted in decreased actomyosin fibers and increased vascular permeability.

We next assessed VEC barrier function *in-vivo* using intravital two-photon microscopy, by quantifying fluorescence intensity in *Sema3f^-/-^* ear blood vessels after FITC-dextran injection. Fluorescence intensity was decreased over time in blood vessels (ANOVA p=0.03, n=3 each) (Figure 4C), and increased in the peri-vascular space of *Sema3f^-/-^* mice (*WT*: 1.06±0.03; *Sema3f^-/-^*: 2.88±0.11, fold change 20min/0min, n=3 each, p<0.0001) (Figure 4D), suggesting increased vascular permeability. Immunohistological staining of *Sema3f^-/-^* aortae with VE-cadherin showed disordered VECs (Figure 4E), confirming that disturbed VEC organization and increased endothelial permeability likely contributed to increased monocyte migration in these mice.

The decreased VE-cadherin expression in *Sema3f^-/-^* aortas was confirmed by Western blot (*WT*: 1.8±0.07; *Sema3f^-/-^*: 1.4±0.03, fold change, n=4 each, p=0.029) (Figure 4F). Previous studies established that the PI3K/mTOR pathway regulates VE-cadherin expression (28). Phosphorylated mTOR levels were decreased in *Sema3f^-/-^* aorta (*WT*: 1.60±0.01; *Sema3f^-/-^*: 1.26±0.10, fold change normalized to total mTOR, n=4 each, p=0.029) (Figure 4G), suggesting that decreased mTOR activation likely contributed to the decreased VE-cadherin expression and increased vascular permeability in *Sema3f^-/-^* aorta. Since aorta consist of several cell types, we determined phosphorylated mTOR levels in isolated aortic VECs and found it decreased in *Sema3f^-/-^* mice (*Sema3f^+/+^*: 1.12±0.11; *Sema3f^-/-^*: 0.56±0.008, fold change normalized to total mTOR, n=3 each, ANOVA p=0.002), which was reversed upon treatment of *Sema3f^-/-^* VECs with SEMA3F protein (*Sema3f^-/-^*: 0.56±0.008; *Sema3f^-/-^* + SEMA3F: 1.55±0.16, fold change normalized to total mTOR, n=3 each, ANOVA p=0.002) (Figure 4H).

### SEMA3F reduces vascular smooth muscle cell proliferation and migration

Vascular smooth muscle cells (VSMCs) play a critical role in atherogenesis, and pathological VSMC proliferation and migration is a major factor in neointimal lesion formation (29). It is estimated that up to 70% of the mass of atherosclerotic lesions can be VSMCs or VSMC-derived (30). We found decreased lesional VSMCs in *Ldlr^-/-^* mice administered SEMA3F (Figure 1C), and increased lesional VSMCs in *Sema3f^-/-^* mice (Figure 2C), suggesting that reduced VSMC proliferation and/or migration may contribute to SEMA3F-mediated protection against neointima formation. Indeed, SEMA3F decreased primary human coronary artery smooth muscle cell (HCASMC) proliferation (ANOVA p=0.0008, Supplemental Figure 13). In addition, HCASMC migration was decreased by ∼40% at 4 h (PBS: 24.1±2.6, n=9; SEMA3F: 14.6±1.8, n=10, % gap reduction, ANOVA p<0.0001, Figure 5A), by ∼30% at 8 h (PBS: 46.9±2.8, n=9; SEMA3F: 33.4±1.9, n=10, % gap reduction, ANOVA p<0.0001, Figure 5A), and by ∼25% at 12 h (PBS: 61.8±2.4, n=9; SEMA3F: 46.8±2.0, n=10, % gap reduction, ANOVA p<0.0001, Figure 5A) in response to SEMA3F treatment.

**Figure 5:**
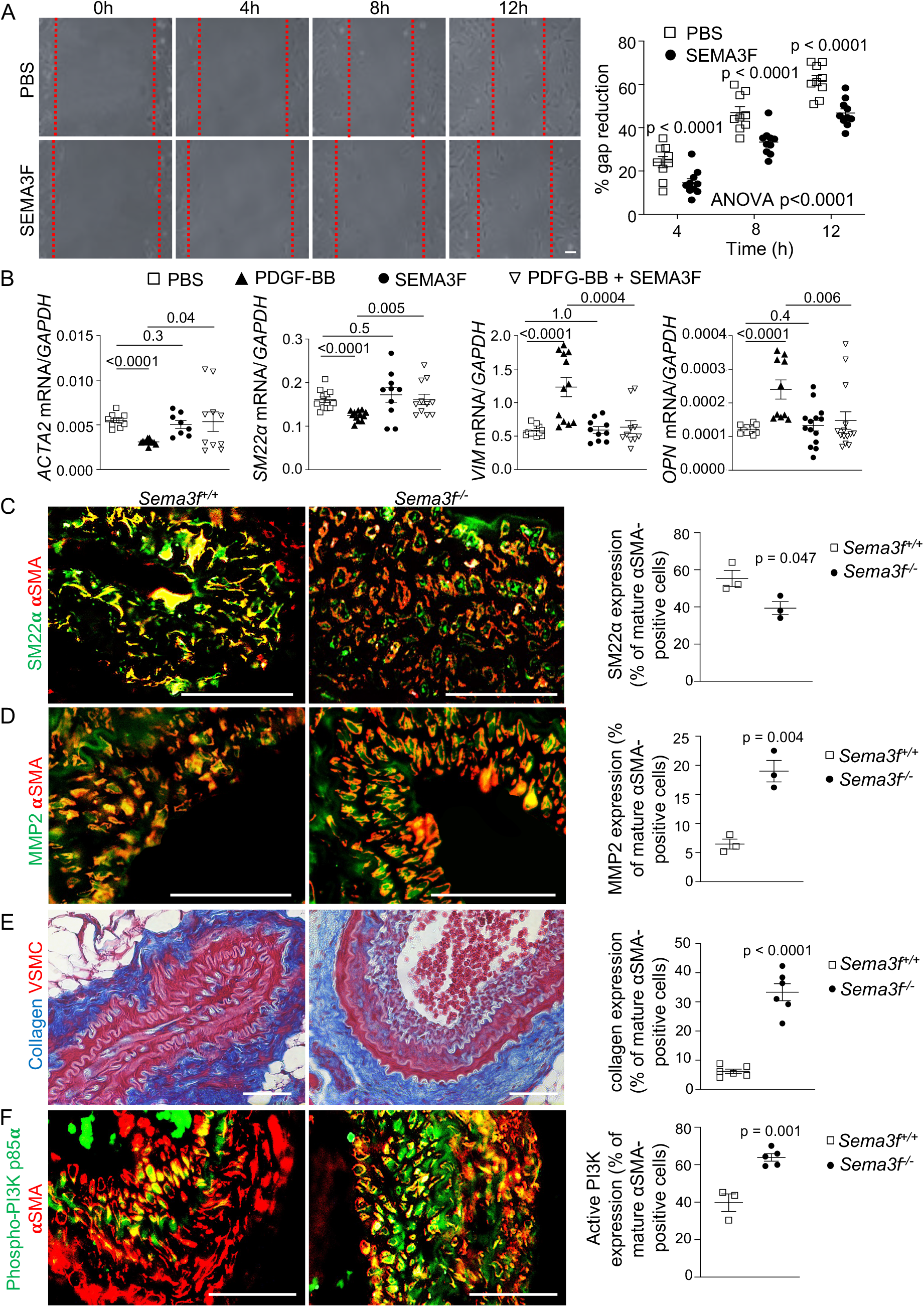
SEMA3F regulates migration, contractile phenotype and PI3K activation in vascular smooth muscle cells. **(A)** SEMA3F reduces primary human coronary artery smooth muscle cell migration. **(B)** SEMA3F maintains normal vascular smooth muscle cell contractile phenotype, and SEMA3F treatment rescues the PDGF-BB-induced pro-atherosclerotic synthetic phenotype in VSMCs as assessed by quantifying expression of the contractile smooth muscle markers Smooth muscle actin alpha 2 (*ACAT2*) and Transgelin (*SM22α*), and the synthetic phenotype markers (Vimentin) *VIM* and Osteopontin (*OPN*). **(C)** The number of VSMCs expressing SM22α, a contractile marker (green), as a percentage of mature VSMCs expressing αSMA (red) are decreased in plaques from *Sema3f^-/-^* mice. n=3 each. **(D)** MMP2 positive cells (green), as a percentage of αSMA positive VSMCs (red) are increased in *Sema3f^-/-^* plaques. n=3 each. **(E)** Increased plaque collagen area assessed by Gomori staining, as a percent of total vessel area in *Sema3f^-/-^* mice. n=6 each. Collagen = blue, VSMC = red. **(F)** Cells expressing active PI3K, assessed by phospho-PI3K p85α (Tyr607) staining (green), as a percentage of αSMA positive VSMCs (red) are increased in *Sema3f^-/-^* plaques. n=3-5. Abbreviations are: *Sema3f^+/+^*, *wild-type* mice; *Sema3f^-/-^*, *Sema3f* knockout mice; PDGF-BB, platelet-derived growth factor BB; MMP2, matrix metalloproteinase 2. Values are mean±SEM. Data in (A) was analyzed using Two-way ANOVA followed by Tukey’s multiple comparison tests. Data for *ACAT2* and *SM22*α in (B) were analyzed using Student’s *t*-tests, and data for *VIM* and *OPN* in (B) were not normally distributed and analyzed using Mann-Whitney *U*-tests. Data in (C) to (F) were assessed using Student’s *t*-tests. Scale bar in (A): 100 µm, (C) to (F): 25 µm.

### Decreased contractile vascular smooth muscle cell markers and increased extra-cellular matrix in *Sema3f^-/-^* lesions

VSMCs express contractile markers, whose expression is reduced during atherosclerosis and neointima formation, leading to switching of VSMCs from contractile to a more pro-atherogenic synthetic phenotype associated with increased cell proliferation, migration and secretion of extracellular matrix proteins (29). Thus, we quantified expression of the canonical contractile genes Smooth muscle actin alpha 2 (*ACTA2*) and Transgelin (*TAGLN* or *SM22α*), and the well-characterized markers for synthetic VSMCs, Vimentin (*VIM*) and Osteopontin (*OPN*) (31, 32), in SEMA3F-treated HCASMCs. Platelet-derived growth factor-BB (PDGF-BB) switches VSMCs from contractile to the synthetic phenotype (33). Thus, we utilized PDGF-BB as a positive control, and found that PDGF-BB treatment reduced both *ACTA2* and *SM22α* expression, and increased *VIM* and *OPN* expression, as expected (Figure 5B). In contrast, SEMA3F treatment did not affect *ACTA2* (PBS: 0.0055±0.0002, n=10; SEMA3F: 0.005±0.0004, n=8, relative fold change/GAPDH, p=0.3, Figure 5B), *SM22α* (PBS: 0.16±0.006, n=12; SEMA3F: 0.17±0.02, n=10, relative fold change/GAPDH, p=0.5, Figure 5B), *VIM* (PBS: 0.58±0.02, n=12; SEMA3F: 0.59±0.05, n=10, relative fold change/GAPDH, p=1.0, Figure 5B) or *OPN* expression (PBS: 0.00012±0.000004, n=11; SEMA3F: 0.00013±0.00002, n=14, relative fold change/GAPDH, p=0.4, Figure 5B), indicating that SEMA3F maintains the normal contractile phenotype of HCASMCs. Treating HCASMCs with both PDGF-BB and SEMA3F rescued the synthetic phenotype caused by PDGF-BB (Figure 5B), indicating that SEMA3F reversed the pro-atherogenic effects of PDGF-BB on HCASMCs.

To determine if SEMA3F contributed to VSMC phenotype switching in vivo in the plaque, we quantified VSMCs expressing SM22α, which were decreased in lesions from *Sema3f^-/-^* mice that had undergone carotid partial ligation (*Sema3f^+/+^*: 55.3±4.4; *Sema3f^-/-^*: 39.3±3.5, n=3 each, % of mature αSMA-positive VSMCs, p=0.047, Figure 5C), suggesting that the absence of SEMA3F decreased contractile VSMCs in vivo. These data suggest that SEMA3F maintains the normal contractile phenotype of VSMCs, likely contributing to SEMA3F-mediated atheroprotection.

VSMC switching to a synthetic phenotype during neointimal hyperplasia is characterized by the secretion of extracellular matrix and collagen (31, 34). Lesional MMP2 expression (*Sema3f^+/+^*: 6.5±0.9; *Sema3f^-/-^*: 19.0±1.9, n=3 each, MMP2 expression (% of mature αSMA-positive cells), p=0.004, Figure 5D) and collagen content (*Sema3f^+/+^*: 6.2±0.7; *Sema3f^-/-^*: 33.3±2.9, n=6 each, collagen positive area (% of vessel area), p<0.0001, Figure 5E) were increased in *Sema3f^-/-^* mice with partial carotid ligation, again suggesting that reduced VSMC phenotype switching contributes to the atheroprotective effects of SEMA3F.

### SEMA3F modulates PI3K levels in lesional VSMCs

PI3K regulates VSMC proliferation, migration and phenotype switching (35), and active PI3K levels were increased in *Sema3f^-/-^* VECs. In addition, SEMA3F rescued the pro-atherogenic synthetic VSMC phenotype induced by PDGF-BB (Figure 5B). PDGF-BB induces the synthetic phenotype in VSMCs via an extracellular signal-regulated kinase (ERK) and phosphatidylinositol 3-kinase/Akt (PI3K/AKT)-mediated pathway, resulting in reduced *ACTA2* and *SM22α* activity (36), suggesting that PI3K activity maybe increased in *Sema3f^-/-^* VSMCs. Indeed, phospho-PI3K p85α (Tyr607), a marker of PI3K activity (37), was increased in vivo in lesional αSMA-positive *Sema3f^-/-^* VSMCs (*Sema3f^+/+^*: 39.8±4.7, n=3;*Sema3f^-/-^*: 63.9±2.0, n=5; phospho-PI3K expression (% of mature SMCs), p=0.001, Figure 5F), suggesting that SEMA3F-mediated modulation of PI3K activity may contribute to the pro-atherogenic VSMC phenotypes in *Sema3f^-/-^* mice.

### SEMA3F administration reduces neointimal smooth muscle cells in mice

We next assessed if increased SEMA3F led to decreased neointima formation in hyperlipidemic *Ldlr^-/-^* mice subjected to wire injury-mediated endothelial denudation of the common carotid artery, an established model of VSMC proliferation and migration-induced neointima formation (38). In addition, insufficient re-endothelialization after vascular injury delays vascular healing and increases neointimal lesions (39), making this an ideal model to confirm the impact of SEMA3F on re-endothelialization and plaque stability. SEMA3F administration reduced neointimal plaque area in this model (Control: 53.3±3.5; SEMA3F: 21.4±3.1; % intimal plaque area/vessel area, n=7 each, p=0.0006) (Figure 6A), suggesting a beneficial impact of SEMA3F on VSMCs. In line with this, smooth muscle cell numbers were decreased in intimal lesions of SEMA3F-treated mice (Control: 114.0±14.2; SEMA3F: 64.7±11.5; n=7 each, p=0.03) (Figure 6B). SEMA3F administration also accelerated re-endothelialization of the wire-injured carotid by ∼40% (Control: 47.4±4.2; SEMA3F: 67.5±3.4; % endothelial area/vessel area, n=7 each, p=0.004) (Figure 6C), suggesting increased plaque stability. Intra-plaque macrophage numbers were also decreased (Control: 60.7±8.8; SEMA3F: 15.0±5.1; n=7 each, p=0.0006) (Figure 6D). These data show, in a model of increased VSMC proliferation and migration-induced neointima formation, that SEMA3F decreases neo-intimal plaque area, in part via decreasing pro-atherogenic VSMC phenotypes, as well as increasing re-endothelialization.

**Figure 6:**
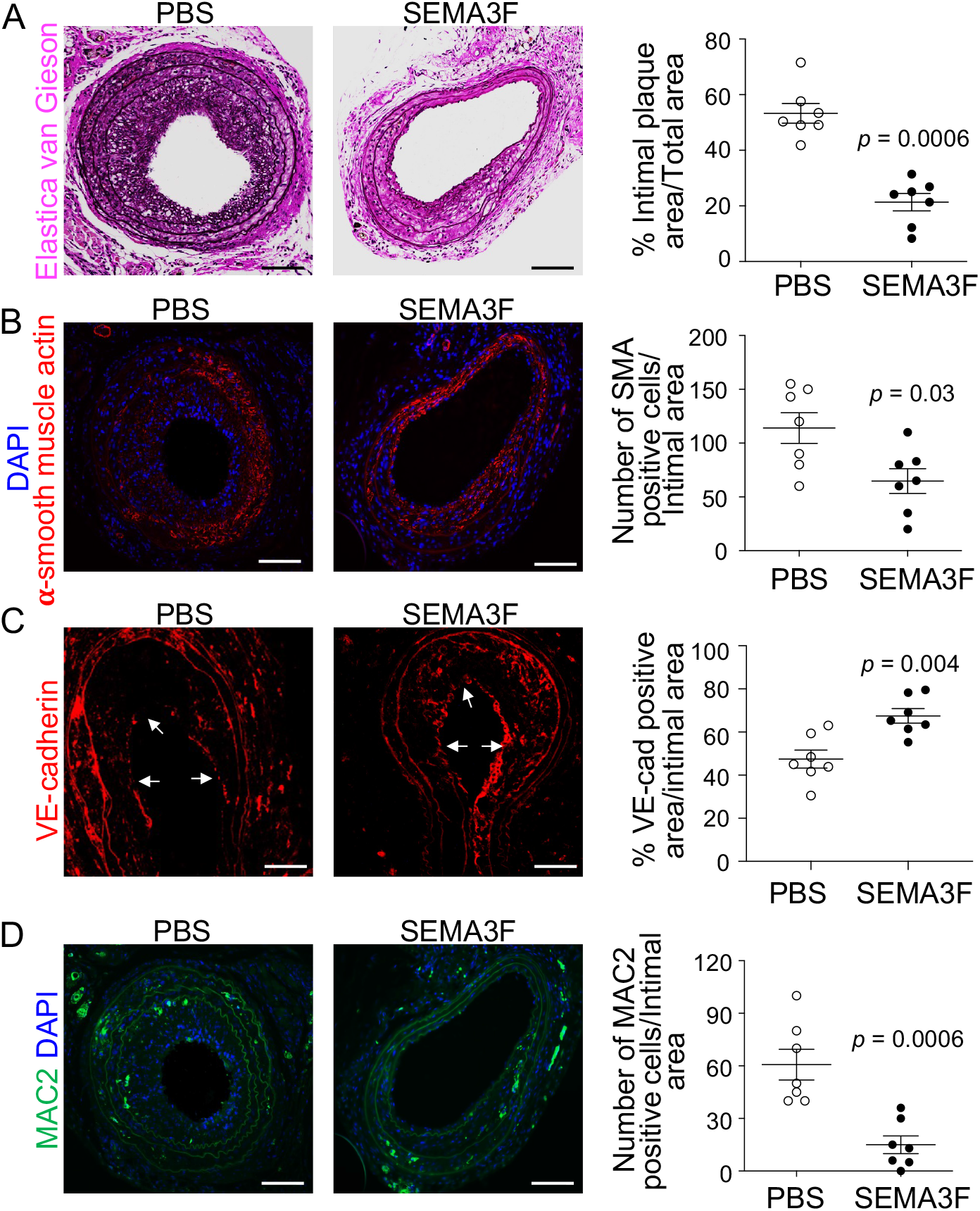
SEMA3F administration reduces intimal lesion area and increases re-endothelialization in a carotid wire injury model of smooth muscle cell proliferation and migration-induced neointima formation. **(A)** Decreased carotid neointimal lesion area in elastica von Gieson-stained carotid sections in SEMA3F-treated *Ldlr^-/-^* mice that underwent common carotid artery endothelial denudation. **(B)** Decreased intimal SMA positive (red) cell numbers in SEMA3F-treated mice. DAPI (blue). **(C)** SEMA3F increased intimal lesion VE-cadherin positive area (red) and accelerated re-endothelialization of the wire-injured carotid endothelium. **(D)** Decreased MAC2 positive macrophages (green) in the neointima. Groups abbreviated as: PBS, carotid wire injured, western-type diet-fed *Ldlr^-/-^* mice, injected with PBS; SEMA3F, carotid wire injured, western-type diet-fed *Ldlr^-/-^* mice, injected with SEMA3F protein. Values represent mean±SEM, 6 sections/mouse, n=7/group. Data were not normally distributed and were analyzed using the Mann-Whitney *U*-test. Scale bar in (A), (B), and (D): 50 µm, (C): 25 µm.

To confirm that pro-atherogenic VSMC phenotypes were indeed decreased, we next determined phenotype switching of lesional VSMCs in the wire injured SEMA3F administered mice, and found that SM22α-expressing anti-atherogenic contractile VSMCs were increased (Control: 43.3±4.8; SEMA3F: 61.0±2.1, n=3 each, % of mature αSMA-positive VSMCs, p=0.03) (Figure 7A). Both MMP2 expression (Control: 41.2±4.2; SEMA3F: 17.2±2.1, n=3 each, MMP2-positive cells (% of mature αSMA-positive cells), p=0.007) (Figure 7B), and collagen content (Control: 40.4±4.2; SEMA3F: 27.0±1.3, n=6 each, collagen positive area (% of vessel area), p=0.01) (Figure 7C), were decreased in plaques in SEMA3F-treated mice with endothelial denudation, again suggesting that reduced pro-atherogenic VSMC phenotype switching contributes to the atheroprotective effects of SEMA3F.

**Figure 7:**
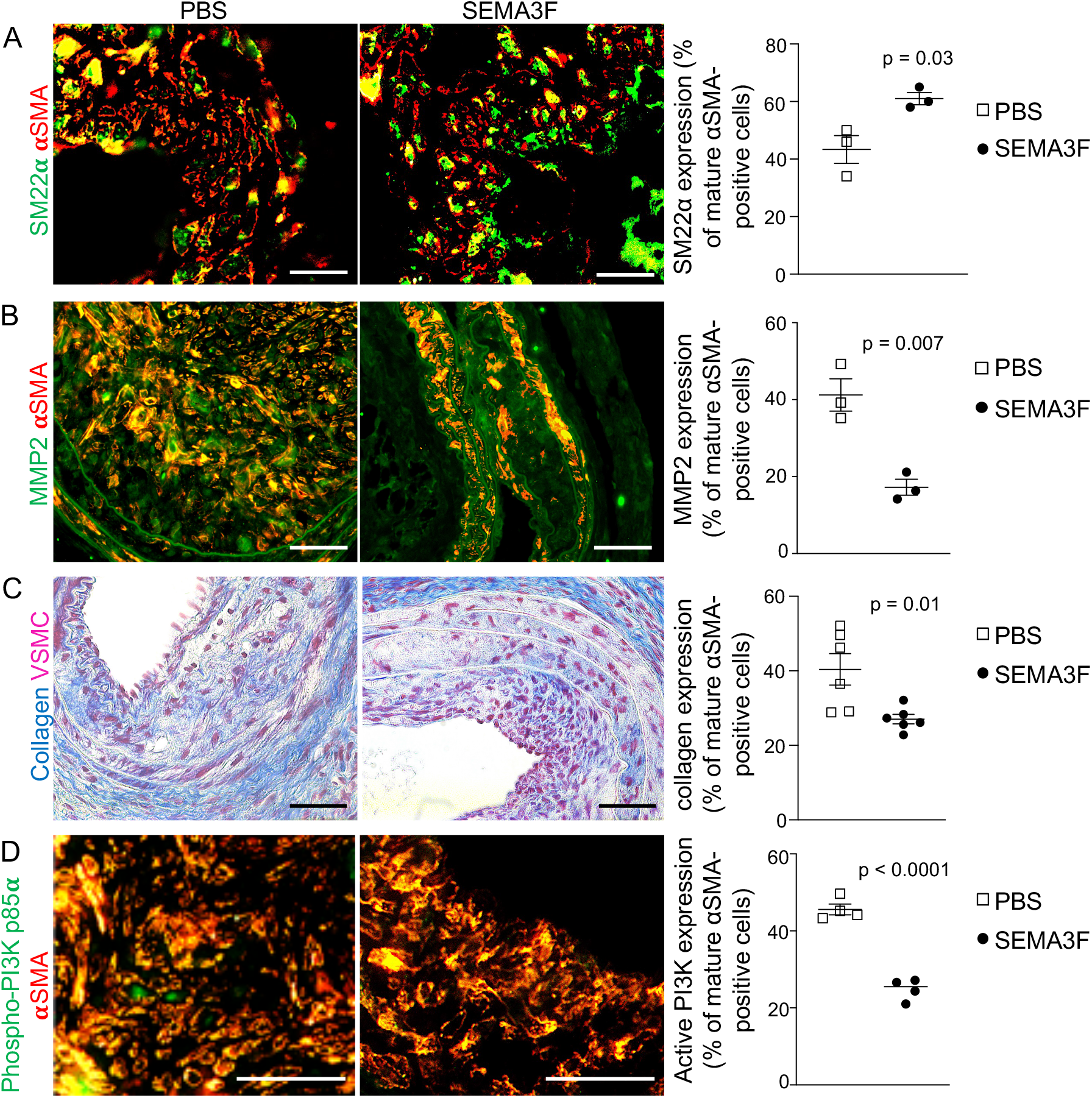
SEMA3F regulates phenotype switching and PI3K activity in vascular smooth muscle cells. **(A)** The number of VSMCs expressing SM22α, a contractile marker (green), as a percentage of mature VSMCs expressing αSMA (red) are increased in neointimal lesions from SEMA3F-treated mice. n=3 each. **(B)** MMP2 positive cells (green), as a percentage of αSMA positive VSMCs (red) are decreased in neointimal lesions from SEMA3F-treated mice. n=3 each. **(C)** Decreased plaque collagen area assessed by Gomori staining, as a percent of total vessel area in neointimal lesions from SEMA3F-treated mice. n=6 each. Collagen = blue, VSMC = red. **(D)** Cells expressing active PI3K, assessed by phospho-PI3K p85α (Tyr607) staining (green), as a percentage of αSMA positive VSMCs (red) are decreased in neointimal lesions from SEMA3F-treated mice. n=4 each. Abbreviations are: MMP2, matrix metalloproteinase 2. Values are mean*±*SEM. Data in (A) to (D) were normally distributed and assessed using Student’s *t*-tests. Scale bar in (A) to (D): 25 µm.

To determine if reduced VSMC PI3K activity in the presence of increased SEMA3F may have contributed to the decrease in pro-atherogenic VSMC phenotypes, we co-stained the neo-intimal plaques with αSMA antibody, to identify mature contractile VSMCs, and phospho-PI3K p85α (Tyr607) antibody, to identify activated PI3K, and found decreased active PI3K in VSMCs of SEMA3F-treated mice (Control: 45.6±1.4; SEMA3F: 24.8±1.4; n=4 each, plaque phospho-PI3K p85α expression (% of mature SMCs), p<0.0001) (Figure 7D). Together, our data imply that SEMA3F protects against neointimal plaque formation via increasing plaque stability and by decreasing the pro-atherogenic phenotypes of lesional VECs and VSMCs, and that SEMA3F-mediated inhibition of the PI3K pathway may contribute to these atheroprotective phenotypes.

## Discussion

This study establishes a causal and protective role for SEMA3F in atherogenesis. Our results extend and provide complementary functional validation to our previously published findings from genome-wide association studies where the semaphorin signaling pathway was significantly associated with coronary artery disease in humans (4). Although, secreted class 3 semaphorins have been implicated in several pathological processes including regulation of immune-cell trafficking (40), inhibition of endothelial cell migration (41), increased vascular permeability (42, 43), and response to oxidized phospholipids in macrophages (44), they have mostly been studied in the context of cancer and tumor angiogenesis (45). SEMA3F has also been shown to reduce monocyte transmigration in venous endothelial cells (7). However, venous cells are not an accurate model for atherosclerosis (24), and a role for SEMA3F in atherogenesis has not been described. Thus, our findings provide the first evidence for a causal role of SEMA3F in atherosclerotic disease. Therapies targeting upregulation and/or activation of SEMA3F signaling pathways may thus prove useful in reducing neointimal plaque formation.

We find that SEMA3F reduces plaque formation through its impact on vascular endothelial and smooth muscle cells. However, we found SEMA3F to also be highly expressed in macrophages. Whether macrophage SEMA3F plays a role in reducing atherosclerosis is unknown and needs further investigation. SEMA3F reduced the transmigration of TNFα-stimulated monocytes (7), suggesting a direct effect of SEMA3F on these cells.

A common mechanism, regulation of PI3K activity by SEMA3F, likely underlies the effects of SEMA3F in VECs and VSMCs, and in both cell types, the loss of SEMA3F led to increased PI3K activity. NF-κB is activated by PI3K/AKT via IKK (19), and we found that PI3K inhibition reduced the NF-κB activation in the absence of SEMA3F. SEMA3F binding to its receptors NRP2/PlexinA1 has been suggested to reduce PI3K activity (20). Together these data suggest an upstream mechanism by which SEMA3F might regulate NF-κB activity, thus decreasing expression of the adhesion molecules ICAM and VCAM1 at the cell surface, leading to decreased monocyte adhesion.

The increase in VEC membrane permeability in the absence of SEMA3F may also be modulated, at least in part, by PI3K-mediated mechanisms. PI3K directly regulates endothelial cell barrier function via modulating the tyrosine phosphorylation of VE-cadherin, a key protein in endothelial cell adherens junctions, thus increasing barrier permeability by altering the actin cytoskeleton/adherens junction complex (46). As well, blocking of mTOR reduces VE-cadherin expression (28), thus increasing VEC membrane permeability. We found increased cell-cell adherens junction thickness upon SEMA3F treatment, and VE-cadherin expression was decreased in the absence of SEMA3F. This was associated with decreased mTOR phosphorylation in aortas and isolated aortic VECs from *Sema3f^-/-^* mice, suggesting that reduced mTOR activity may underlie the decrease in VE-cadherin expression. PI3K regulates mTOR phosphorylation (47), and an in vitro study using glioblastoma cells found that SEMA3F reduced mTOR phosphorylation (20), whereas we found reduced mTOR phosphorylation in the absence of SEMA3F. The reasons for this difference are unclear. However, the previous study used a glioblastoma cell line, whereas we assessed mTOR phosphorylation in aortas and aortic VECs isolated from *Sema3f^-/-^* mice. SEMA3F may play distinct roles in tumorigenesis as opposed to atherosclerosis. In addition, our data may reflect the entire physiological milieu, since they were generated using in vivo and ex vivo studies, and not cell lines. Finally, mTOR associates with RAPTOR to form the mTORC1 functional complex, and associates with RICTOR in forming the mTORC2 complex (48). mTORC2, in concert with PI3K and AKT, can regulate mTORC1, and PI3K also regulates mTORC2 (20), suggesting a complex interplay which may be tissue and function dependent.

In VSMCs, PI3K activation reduces contractile gene expression (49). As well, PI3K activation increases VSMC proliferation and migration (35). We found increased active PI3K in VSMCs of the *Sema3f^-/-^* mice, suggesting that increased PI3K activation in the absence of SEMA3F likely contributed to the pro-atherogenic phenotypes in the *Sema3f^-/-^* VSMCs.

Together, our findings establish SEMA3F in the vascular endothelium and vascular smooth muscle cells as an atheroprotective agent. This atheroprotective role of SEMA3F in the vascular endothelium may share functional similarities with its previously described role as an inhibitor of tumor angiogenesis and a negative regulator of lymphatic endothelial proliferation (50), as well as a regulator of monocyte migration (9). To our knowledge, a role for SEMA3F in decreasing atherosclerosis has thus far not been described.

In addition to the upregulation of the ‘cell adhesion molecules, CAMs’ pathway, using whole-genome expression profiling, we identified that the ‘ECM receptor interaction’ pathway was downregulated in *Sema3f^-/-^* VECs. This finding suggests an additional role of SEMA3F in ‘cell-matrix interactions’, and underscores the need for further investigation into the role of endothelial cell SEMA3F in atherosclerosis.

We find that SEMA3F reduces atherosclerosis progression by reducing monocyte adhesion to the vascular endothelium. SEMA3F also maintains vascular endothelial membrane integrity, thus reducing vascular permeability and preventing monocyte transmigration across the endothelial layers, resulting in an overall reduction in atherosclerosis. SEMA3F also reduces the atherogenic increases in VSMC migration proliferation and phenotype switching. These findings offer a more in-depth view of the complexity of semaphorin biology in vascular dysfunction, and advance SEMA3F as a candidate therapeutic target for atheroprotection.

## Methods

### Data Availability

All data generated or analyzed during this study are included in the manuscript and supporting files. Source data are available for all figures and supplemental figures. Raw data for the RNASeq experiment are available at Dryad (doi:10.5061/dryad.8w9ghx3pg).

### Mouse experiments

All experiments were approved by the Biomedical Sciences Institute Singapore Institutional Animal Care Committee. Mice were maintained on a 12 hr dark-light cycle with *ad libitum* access to water and food, which was standard laboratory diet (1324_modified, Altromin GmbH & Co.) unless stated otherwise. All mice were on the *C57BL/6J* background. All experiments were performed on male mice, to reduce variability. All mice were randomly assigned to cohorts, and all experimental data were analyzed blinded.

### Genetic association analysis and transcriptomic data mining

To identify novel associations between established biological mechanisms and CAD, we performed a 2-stage pathway-based gene-set enrichment analysis of 16 genome-wide association study (GWAS) data sets for CAD (available through the CARDIoGRAM consortium) (4), using i-GSEA4GWAS (52), and by querying the Reactome pathway database. From a meta-analyzed discovery cohort of 7 CAD GWAS data sets (9,889 cases/11,089 controls), nominally significant pathways were tested for replication in a meta-analysis of 9 additional studies (15,502 cases and 55,730 controls). To examine semaphorin gene expression levels in human atherosclerosis-relevant samples, we screened the Gene Expression Omnibus (GEO) (6) for human studies for the keywords ‘macrophages’, ‘vascular endothelial cells’, ‘vascular smooth muscle cells’, and ‘atherosclerotic plaques’. Eighteen microarray studies (Affymetrix and Illumina platforms), encompassing 570 samples were analyzed. To enable comparisons between diverse GEO datasets, the expression values from each study were converted into quintiles with Q1 representing the upper 20% and Q5 the bottom 20% of all expression values.

### Western-type diet-fed *Ldlr^-/-^* model of atherosclerosis

Eighteen male *Ldlr^-/-^* mice (*C57BL/6J*) on a 12h light-dark cycle were maintained on standard laboratory diet (1324_modified, Altromin GmbH & Co.) until 12 weeks of age, followed by 10 week western-type diet feeding (WTD; D12079B, Research Diets). Half the mice (assigned randomly) were injected intra-peritoneally (i.p) with 1xPBS, and the other half with 100 ng/kg of purified SEMA3F protein (3237-S3, R&D Systems), thrice per week, for 10 weeks, in conjunction with WTD feeding. The mice had free access to food and water except during a 4-5 h fast prior to blood sampling. Mice were anesthetized at 22 weeks (100 mg/kg ketamine hydrochloride/10 mg/kg xylazine i.p.), bled retro-orbitally, perfused transcardially with 1xPBS, and the aortic arch and thoracic aorta were isolated, and the surrounding fat tissue removed for en face analyses. Other aortic arch and thoracic aorta were snap frozen for RNA and western blot analyses. The hearts were fixed in 4% paraformaldehyde (Sigma) and embedded in paraffin.

### Atherosclerotic lesion analyses

Serial cross sections (5μm), taken throughout the entire aortic root (9), were stained with hematoxylin-phloxine-saffron and atherosclerotic lesion area was analyzed (6 cross-sections/mouse). Aperio ImageScope 12.1 (Leica Biosystems) and ImageJ were used for quantification of lesion number, area and severity according to the American Heart Association classification (9). Briefly, mild lesions were described as lesions containing a single or multiple layers of foam cells with fibrous caps. Advanced lesions were defined as lesions showing well-formed necrotic cores and cholesterol clefts with overlying fibrous cap and some degree of macrophage invasion with/without intraplaque hemorrhage. MAC-3 (550292, BD-Pharmingen), MAC-2 (CL8942AP, Cedarlane), *α*-SMA (M085129, DAKO) and VE-cadherin (144347, US Biological) antibodies were used to determine lesional macrophage, smooth muscle actin, and endothelial cell content in the intimal lesion area. SEMA3F antibody (SAB2107196, Sigma) was used to co-stain with MAC2, *α*-SMA, and VE-cadherin to evaluate co-localization of SEMA3F in macrophages, smooth muscle and endothelial cells, respectively.

### *En-fac*e staining of whole aorta

Aorta were isolated, cleaned of perivascular adipose tissue, opened longitudinally, pinned on dissection wax to expose the intimal surface, rinsed 3 times with 1x PBS, fixed with 70% ethanol for 5 min at room temperature, and stained with Oil Red O (Sigma-Aldrich) for 20 min at room temperature. Aorta were rinsed with water and imaged. ImageJ was used to quantify lesion and total arch area.

### Partial ligation model of atherosclerosis

Four month-old male *Sema3f^-/-^* mice (*C57BL/6J* background) (51) were sedated (2% isoflurane), a ventral midline neck incision was made, and muscle layers separated to expose the left carotid artery (LCA). Three of four branches of the LCA (left external carotid, internal carotid, and occipital arteries) were ligated using 6-0 silk sutures. The superior thyroid artery was left intact, providing the sole source of blood circulation. The incision was closed, and the mice kept on heating pads until conscious. Five weeks after injury, mice were perfused with 4% paraformaldehyde. The injured carotid arteries were isolated in 4% paraformaldehyde, dehydrated, and paraffin embedded. Serial 5μm transverse sections (6 sections/mouse, 75μm apart) were collected within 0 to 480μm from the bifurcation and stained with Elastica-van Gieson (EVG) (Sigma). Areas of lumen, intima and media were measured by planimetry using Aperio ImageScope 12.1 (Leica Biosystems). For each mouse, data from the 6 sections were averaged to represent lesion formation along this standardized distance. The intimal lesion area was calculated as a percentage of total vessel area.

### Plasma lipoprotein quantification

Plasma total cholesterol was quantified using the Infinity Cholesterol Reagent (Thermo-Fisher Scientific). HDL cholesterol was quantified in the supernatant after ApoB-containing particles were precipitated using 20% polyethylene glycol (PEG) in 0.2M glycine, using the Infinity Cholesterol reagent. Non-HDL cholesterol was calculated by subtracting HDL cholesterol from total cholesterol per mouse. Plasma triglycerides were measured using the LabAssay^TM^ Triglyceride kit (WAKO, Fujifilm).

### Isolation of mouse primary aortic endothelial cells

Mice were anesthetized at 8 weeks (100mg/kg ketamine hydrochloride/10mg/kg xylazine i.p.) and perfused transcardially with 1xPBS. Thoracic aortas were removed in cold PBS, opened longitudinally, and cut into 1-mm-wide strips, which were placed lumen-face down in 12-well culture plates coated with Matrigel (354234, Corning™). Plates were incubated in a humidified cell culture incubator at 37°C until the Matrigel solidified. EndoGRO-VEGF Complete Culture Media (Sigma) was added to the wells, and the aortic strips cultured for 4 days. Following cell colony formation, the remaining aortic tissue was removed, and cell culture continued until confluent. The cell colonies were trypsinized and transferred to collagen-coated flasks with EndoGRO-VEGF media until confluent. Before experiments, the purity of the endothelial cells was determined using CD31 MicroBeads (Miltenyi Biotec). Briefly, the total cell number was counted, and cells labeled with CD31 MicroBeads in PBS, 0.5% BSA and 2 mM EDTA. The cell suspension was loaded onto a MACS® column (Miltenyi Biotec), which was placed in a magnetic MACS separator. The magnetically labeled CD31^+^ cells were retained within the column, and subsequently eluted via removal of the column from the magnetic field. The number of CD31^+^ cells were >80% of total cells in all experiments.

### Isolation of mouse splenic cells and monocyte purification

The spleen was dissociated from male mice in medium (RPMI 1640/10%FBS/1% pen-strep) and cells passed through a 40 μm cell strainer, washed, and pelleted. Red blood cells (RBCs) were lysed in RBC lysis buffer, washed with cold 1XPBS and filtered through a 70 μm cell strainer. Monocytes were purified using EasySep™ Mouse Monocyte Isolation Kit (19861A, Stemcell Technologies) following manufacturer’s instructions.

### Monocyte adhesion assays

Passage numbers 1 or 2 of isolated VECs were cultured in a modified 3D microfluidic vasculature chip developed previously (53), with channel size of ∼1.5cm length and ∼750μm diameter, and coated with 50μg/mL fibronectin (Sigma-Aldrich). Confluent cell monolayers were dissociated using 0.25% trypsin with 1mM EDTA (Gibco), and the VEC suspension (2 million cells/ml) was seeded into the channel twice to facilitate cell adhesion onto the bottom surface and the top surface of the 3D channel. The chips were then cultured until confluent before performing the monocyte adhesion assays.

In some experiments, VECs were treated with SEMA3F protein (100ng/ml) overnight, or anti-VCAM1 or anti-ICAM1 antibodies (40ng/ml each) for 2 h, following which purified mouse splenic monocytes (15 million/ml) or Cell Tracker (Thermo Fisher Scientific)-labelled THP-1 cells (2 million/ml) were perfused through the microchannel at an inlet wall shear stress of 3 dyne/cm^2^ (atherogenic shear stress ∼1-4 dyne/cm^2^) for 10 minutes using a peristatic pump (P720, Instech Laboratories). The channel was washed using culture media and the number of monocytes adhered to the VECs were imaged using a microscope (Nikon Eclipse Ti) and quantified using ImageJ (National Institutes of Health). The experiments were replicated thrice, with cells from different donor mice.

### RNA sequencing

Total RNA was purified by RNeasy kit (Qiagen). 150 base, paired-end RNA sequencing reads (average read depth >53 million) were generated on Illumina Novaseq (NovogeneAIT Genomics), and aligned to the mouse reference genome (GRCm38) via STAR aligner (version 2.5.1b) with an average mapping rate of 95 percent. Mapped sequencing reads were quantified and assigned to genomic features via the *featureCount* option in R package *Rsubread*. Differential gene expression analysis was conducted in R via *limma* (54). Genes with absolute log2 fold change ≥1.0 and adjusted p-value ≤0.05 were considered as differentially expressed. Transcriptome data was further subjected to pathway enrichment analysis by querying ‘gene-sets’ from the Kyoto Encyclopedia of Genes and Genomes (KEGG) (55) pathway repositories via the PreRankedGSEA tool or Enrichr over-representation analysis tool. Pathways with adjusted p-value ≤0.05 were considered to be significantly enriched.

### Western immunoblotting

Mouse aortic tissues (male mice) were collected in lysis buffer (RIPA, Thermo Fisher Scientific) containing phosphatase inhibitors (5870, Cell Signalling) and protease inhibitors (S8820, Sigma), then homogenized. 20μg of protein was separated on 12% SDS-PAGE gels, and transferred onto 0.4mm PVDF membranes (EMD Millipore). After blocking in 5% skim milk for 1 h, membranes were incubated at 4°C overnight with primary antibodies against SEMA3F (SAB2107196, Sigma), NF-κBp65 (8242, Cell Signaling), VCAM1 (16-1061-82, Invitrogen) and ICAM1 (16-0541-81, Invitrogen), β-actin (4970, Cell Signaling), VE-cadherin (144347, US Biological), Calnexin (2433, Cell Signaling), mTOR and PhosphoSer2448-mTOR (2972 and 5536 respectively, Cell Signaling). Membranes were incubated with secondary antibodies for 1 h at room temperature. All protein bands were visualized using ECL solution on a BioSpectrum Imaging System (UVP). The band intensities were analysed using ImageJ (National Institutes of Health).

### NF-κB immunofluorescence

Mouse primary VECs were cultured in 8-well chamber glass slides (80841, Ibidi) and treated with SEMA3F (100ng/ml) or Wortmannin (1µM, sc-3505) overnight. Cells were fixed with 4 % paraformaldehyde and incubated with 0.1% Triton-X100 (Sigma). After blocking in 3% BSA, cells were incubated at 4°C overnight with primary antibody against NF-κBp65 (8242, Cell Signaling). Nuclei were counterstained with DAPI in Vectashield mounting media (Vector Labs). Images were quantified using ImageJ.

### PI3K activity

The levels of active PI3K were determined by quantifying the ratio of phosphorylated PI3K p85 (phospho-tyrosine 467/199) and total PI3K p85, normalized to cell numbers, using an ELISA kit and following the manufacturer’s protocol (ab207484, Abcam).

### Monocyte migration

Monocyte migration through VECs were performed as previously described (56). Briefly, isolated VECs from male mice were placed on the upper membranes of a 3D-Flow Chamber Device (Expedeon), and incubated in a cell culture incubator for 2 h. Media with 10ng macrophage colony-stimulating factor (M-CSF) was added to the bottom chamber, the device was connected to a peristaltic pump and 5×10^6^ monocytes were flowed into the device at 0.2 ml/min for 24h. Migrated monocytes were collected from the bottom chamber and counted using automated cell counting (Countless II, Thermo Fisher Scientific). The experiments were replicated thrice, with cells from different donor mice.

### F-actin immunofluorescence and confocal microscopy

Cells grown on glass bottom dishes (Ibidi) were fixed with 4% PFA for 10 min then permeabilized for another 10 min with 0.1% Triton X-100. The dishes were blocked for 1 h with 5% bovine serum albumin (BSA) in PBS and incubated with primary antibodies overnight at 4°C. Antibodies used were anti-non-muscle myosin IIA (1:400; M8064, Sigma-Aldrich), anti-vinculin (1:200; V9131, Sigma-Aldrich), anti-VE-cadherin (1:200; 2158; Cell Signaling). To visualize F-Actin and nuclei, cells were stained with Alexa Fluor™ 647 Phalloidin (Thermo Fisher) and Hoechst (Sigma) respectively. Microscopy was carried out using a Yokogawa CSU-W1, Nikon TiE spinning disk confocal microscope.

### Vascular endothelial cell focal adhesion dynamics and cell junction analysis

To examine focal adhesion size, HAEC cells (ATCC PCS-100-011) were seeded at sub-confluent levels on glass bottom dishes (Ibidi) and treated with a 100ng/ml SEMA3F or PBS overnight and probed for vinculin expression. Confocal images were taken at the basal plane and focal adhesions were then analyzed using Focal Adhesions Analysis Server (https://faas.bme.unc.edu) to determine the average area (μm^2^) of vinculin adhesions per cell. To examine endothelial cell-cell junctions, HAEC cells were seeded to confluency on glass bottom dishes and treated with a 100ng/ml SEMA3F or PBS overnight and probed for VE-cadherin. Cell monolayers were imaged at 40x magnification and 10μm Z-stacks were acquired and merged into maximum projections. Cadherin junction width was calculated by the measuring the average width of individual junctions between two cells using ImageJ.

### *In-vivo* analyses of vascular permeability in mice

Anesthetized mice (males) were positioned on the stage of a two-photon microscope (LaVision TriM Scope II (LaVision Biotech) with an upright microscope (Olympus BX51/61 WI) equipped with a motorized LaVision BioTec xyz-stage, a two-photon Chameleon (Coherent) Ti:Sapphire tunable laser (680-1050 nm), and Olyumpus LUMFLN 60x/1.10 water dipping objective and PMT detectors), and their right ear lobe fixed beneath cover slips with a single drop of water. 5mg FITC-dextran (40-kDa; Sigma-Aldrich) in 300 μl of PBS was injected into the left retro-orbital sinus, and images acquired every 5 minutes starting immediately after injection. The blood vessel area and intensity were measured using ImageJ. The experiment was performed in 3 mice per group.

### Vascular smooth muscle cell migration

Primary human coronary artery smooth muscle cells (HCASMC, Thermofisher Scientific) were cultured in 231 culture media (Thermofisher Scientific) supplemented with Smooth Muscle Growth Supplement (SMGS, Thermofisher Scientific) and used below 9 passages. HCASMCs (1×10^5^ cells/well) were seeded in 12 well plates in media without serum in order to inhibit cell proliferation. After 24 h, the cell monolayer was scratched in a straight line using a sterile 200 μL pipet tip. Cells were either stimulated with SEMA3F (100ng/ml) or PBS as a control in DMEM supplemented with 2% FBS. Digital images were captured at various timepoints (Nikon Fi-1 L2 camera), and the width of the wound was measured using ImageJ software. The results were expressed as % reduction in gap width.

### Vascular smooth muscle cell proliferation

HCASMC (1×10^5^ cells/dish) were grown in 231 culture media supplemented with Smooth Muscle Growth Supplement (SMGS, Thermofisher Scientific), either stimulated with SEMA3F (100 ng/mL) or PBS vehicle. Media was changed daily, and cells were counted at different time points (24, 48 and 72 h).

### Vascular smooth muscle cell phenotype switching

HCASMCs were seeded in 6 well plates (3×10^5^ cells/well). Cells were stimulated either with Platelet-derived growth factor (PDGF-BB, 10 ng/ml), SEMA3F protein (100ng/mL), or PDGF-BB (10 ng/mL) + SEMA3F (10 ng/mL) for 36 h. Total RNA was isolated from HCASMCs using the RNeasy kit (Qiagen). cDNA was generated using random primers and Superscript II reverse transcriptase (Life Technologies). Quantitative real-time PCR was performed using SYBR Green PCR Master Mix (Applied Biosystems) in an ABI QuantStudio 6 Flex Real-Time System (Life Technologies). Reactions were performed in technical triplicates using specific primers. Primer sequences are available upon request.

### VSMC Histology and immunochemistry

Early differentiation of VSMCs (SM22*α*, Ab14106, Abcam), MMP2 (Ab110186, Abcam) and phospho-PI3K p85α (Tyr607) (Ab182651, Abcam) were assessed by counting positive-stained cells in three different fields per section and were expressed as the percentage of positive cells of the total mature VSMCs (smooth muscle actin *α*SMA, M 0851 clone 1A4, DAKO). Collagen staining was assessed using Gomori’s 1-step trichrome stain (ab150686, Abcam). Blue-stained collagen content was analyzed with Cell P Software (Olympus) and expressed as a percentage of the whole vessel area.

### Carotid artery wire injury

Male, 10-12 week old *Ldlr^-/-^* mice (*C57BL/6J* background) were obtained from Jackson Laboratory, and fed WTD for 1 week before and 3 weeks after wire injury. For the wire injury, mice were anaesthetized (100mg/kg ketamine hydrochloride/10mg/kg xylazine i.p.) and subjected to endothelial denudation of the left common carotid artery by a 1cm insertion of a flexible 0.36mm guide wire through a transverse arteriotomy of the external carotid artery, as described (57, 58). Purified SEMA3F protein (100ng/kg, 3237-S3, R&D Systems) was administered via i.p injections, thrice per week, starting on the day of wire injury and continuing for 3 weeks. Mice were then sacrificed and perfused with 4% paraformaldehyde. Both carotid arteries were isolated, fixed in 4% paraformaldehyde, dehydrated and embedded in paraffin. Serial 5μm transverse sections (6 sections/mouse, 75μm apart) were collected from 0 to 480μm from the bifurcation, stained using EVG and areas of lumen, neointima (between lumen and internal elastic lamina), and media (between internal and external elastic laminae) were measured by planimetry using Aperio ImageScope 12.1 (Leica Biosystems). For each mouse, data from the 6 sections were averaged to represent lesion formation. Neointimal macrophages and smooth muscle cells were visualized by immunofluorescence staining for MAC2 (M3/38, Cedarlane) and SMA (1A4, Dako), respectively, followed by FITC- or Cy3-conjugated secondary antibodies (Jackson Immuno Research) as described (59). The intimal lesion area was calculated as a percentage of total vessel area.

### Statistical analyses

All data were tested for normality using D’Agostino and Pearson or Shapiro-Wilk test. Non-parametric data were analyzed by Mann-Whitney *U*-test, and parametric data were analyzed by unpaired Student’s *t*-tests. One-way ANOVA was used for multiple group analyses, followed by Tukey’s multiple comparisons test. Two-way ANOVA was used to analyze the vessel leakage. Data are shown as average ± standard error. All statistical analyses, except for RNA sequencing data, were performed using GraphPad 9.0. A nominal p<0.05 was considered significant.

### Study Approval

All experiments were approved by the Biomedical Sciences Institute Singapore Institutional Animal Care Committee.

## Abbreviations

SEMA3F: Semaphorin 3F
PI3K: Phosphoinositide 3-kinase
VEC: vascular endothelial cell
VSMC: vascular smooth muscle cell
VCAM: Vascular Cell Adhesion Molecule 1
ICAM: Intercellular Adhesion Molecule 1
NF-κB: Nuclear Factor Kappa B Subunit 1
WT: wild-type
WTD: western type diet
HCAEC: primary human coronary artery endothelial cell
HCASMC: primary human coronary artery smooth muscle cell

## Author contributions

CR, DC-M, CS, CP, MH, KSM, WL, AJF, PN, EAL, DMC and MKS performed experiments and analyzed the data. AA provided the *Sema3f^-/-^* mice. EAL, SCW, OR, HWH, AB, SG, RRS designed experiments, RRS and SG funded the project, CR, SG and RRS wrote the manuscript, and all authors edited the manuscript.

## Acknowledgements

We are grateful to Roya Soltan for the technical assistance. We also thank Thankiah Sudhaharan at the A*STAR Microscopy Platform for help with the multiphoton microscopy. The Microscopy Platform is funded by A*STAR and by SingaScope, a National Research Foundation Shared Infrastructure Support grant.

## SUPPLEMENTAL MATERIAL

**Supplementary Figure 1:**
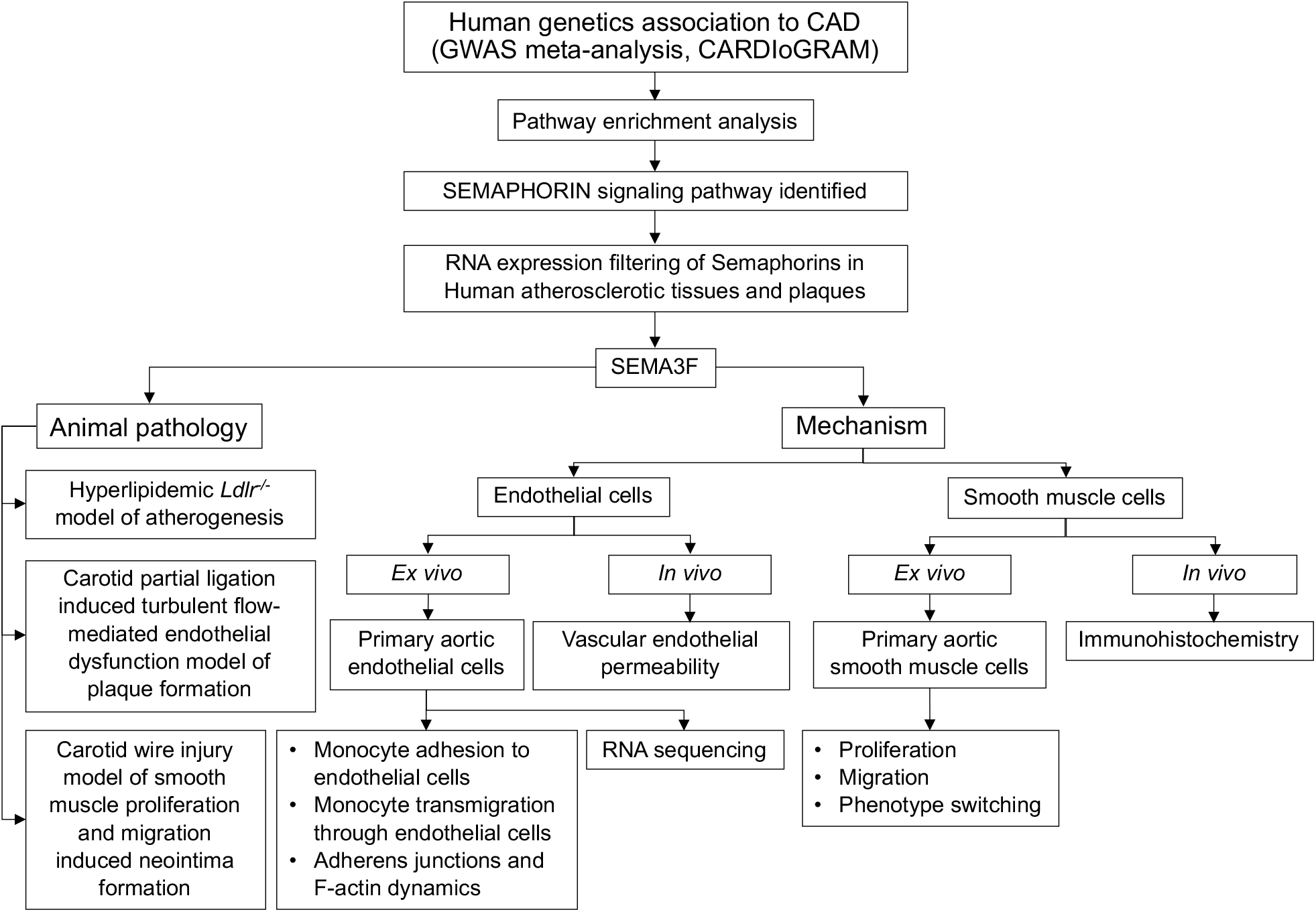
Flow chart of the study approach. Our study expands the clinical relevance of Semaphorins in cardiovascular disease in three important ways. We first identified variants in the Semaphorin signaling genes as significantly associating with cardiovascular disease using the CARDIoGRAM consortium GWAS data (1). In the second step, through knockout and administration studies in mice, the roles of SEMA3F in vascular disease was investigated. Finally, the mechanisms underlying the atheroprotective roles of SEMA3F were identified using *ex-vivo* and *in-vivo* experiments.

**Supplementary Figure 2:**
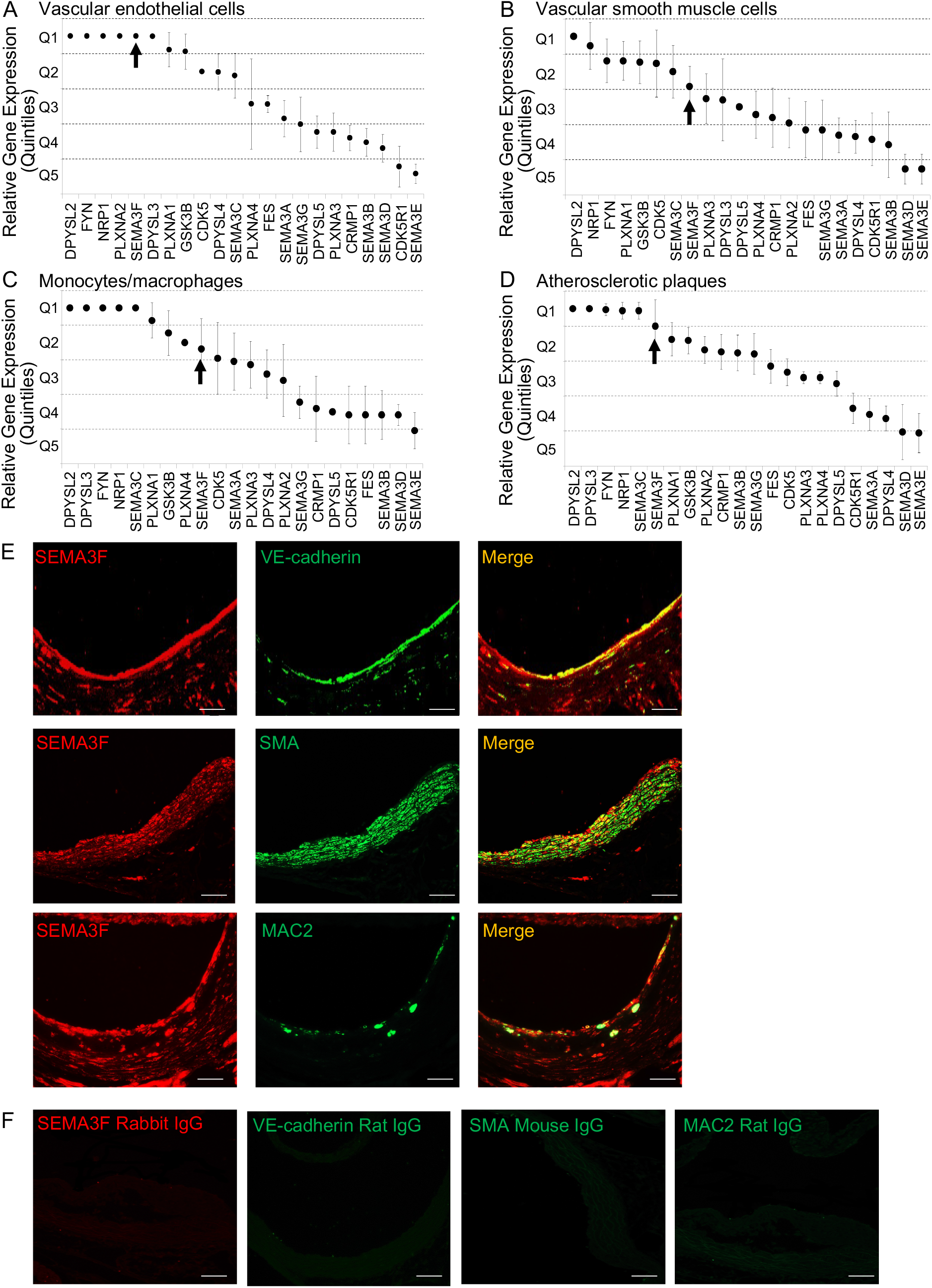
SEMA3F is expressed in human and mouse atherosclerosis-relevant tissues. Expression of genes in the Semaphorin signaling network was assessed in human Gene Expression Omnibus datasets (3), showing high *SEMA3F* expression in **(A)** vascular endothelial cells, **(B)** vascular smooth muscle cells, and **(C)** macrophages. **(D)** *SEMA3F* was also expressed in human atherosclerotic lesions. To allow comparisons between disparate experiments, gene expression levels were converted to quantiles (quintiles) with quintile 1 (Q1) representing the top 20% expressed genes. **(E)** SEMA3F (red) co-localizes with endothelial (VE-cadherin, green), smooth muscle cells (SMA, green), and macrophages (MAC2, green) in the in the aortic sinus of standard laboratory diet-fed *Ldlr^-/-^* mice. **(F)** IgG controls for the antibodies. Scale bar in (D-E): 50 µm.

**Supplementary Figure 3:**
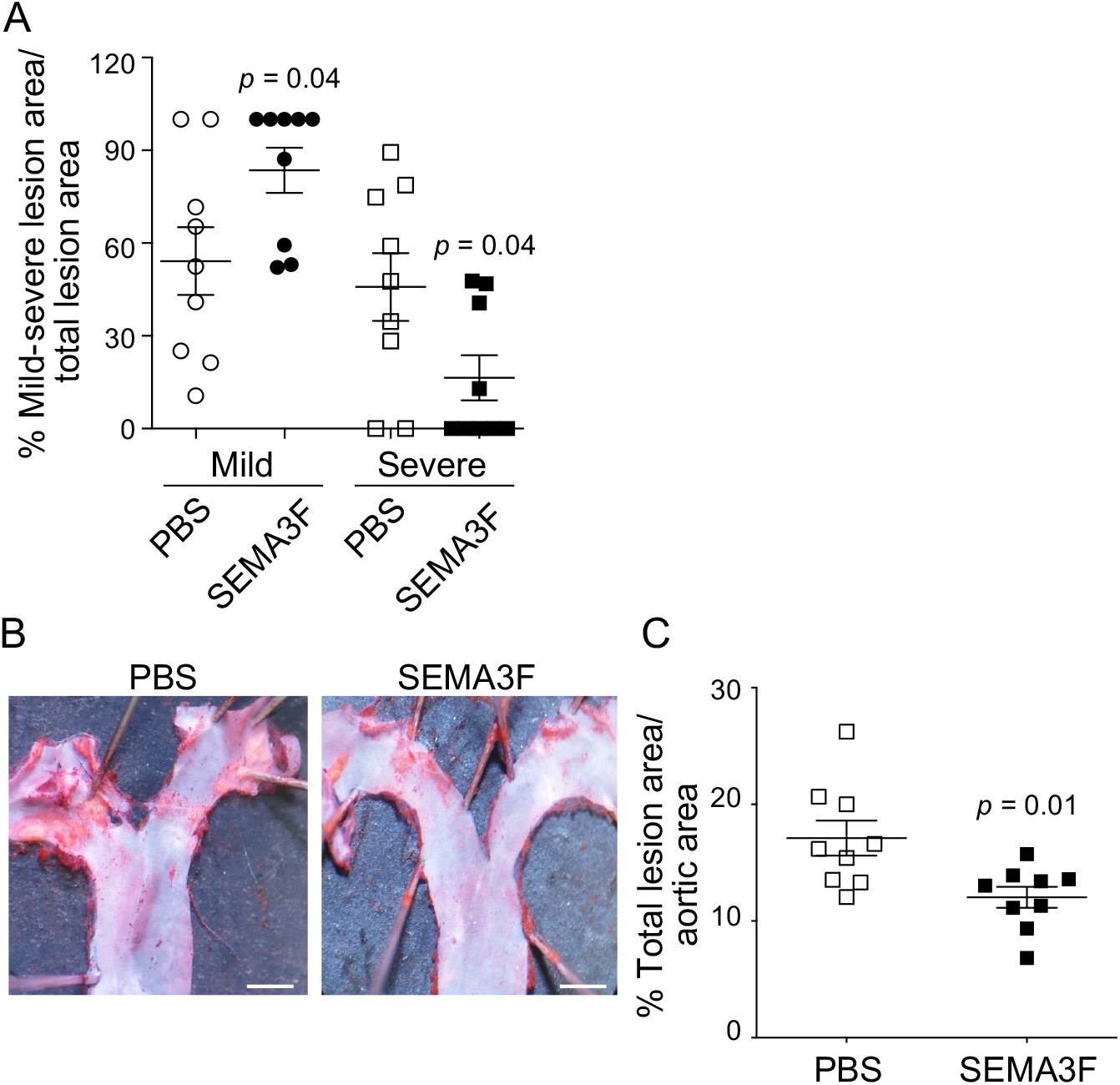
SEMA3F administration reduces atherosclerotic lesion severity and lesion area in *Ldlr^-/-^* mice. **(A)** Significantly lower severe lesion area (types IV and V) and no significant differences in mild lesion area (types I to III) in SEMA3F treated mice. **(B)** Representative photomicrographs of lesion area in *en-face* Oil Red O stained aortic arches. **(C)** Significantly decreased % aortic lesion area (normalized to aortic arch surface area) in the SEMA3F-treated mice. Groups are abbreviated as: PBS, *Ldlr^-/-^* mice fed western-type diet, injected with PBS; SEMA3F, *Ldlr^-/-^* mice fed western-type diet, injected with SEMA3F protein. Values are mean±SEM, 6 serial sections/mouse, n=9/group. Data in (A) and (C) were normally distributed and analyzed using Student’s *t*-tests. Scale bar in (B): 100 µm.

**Supplementary Figure 4:**
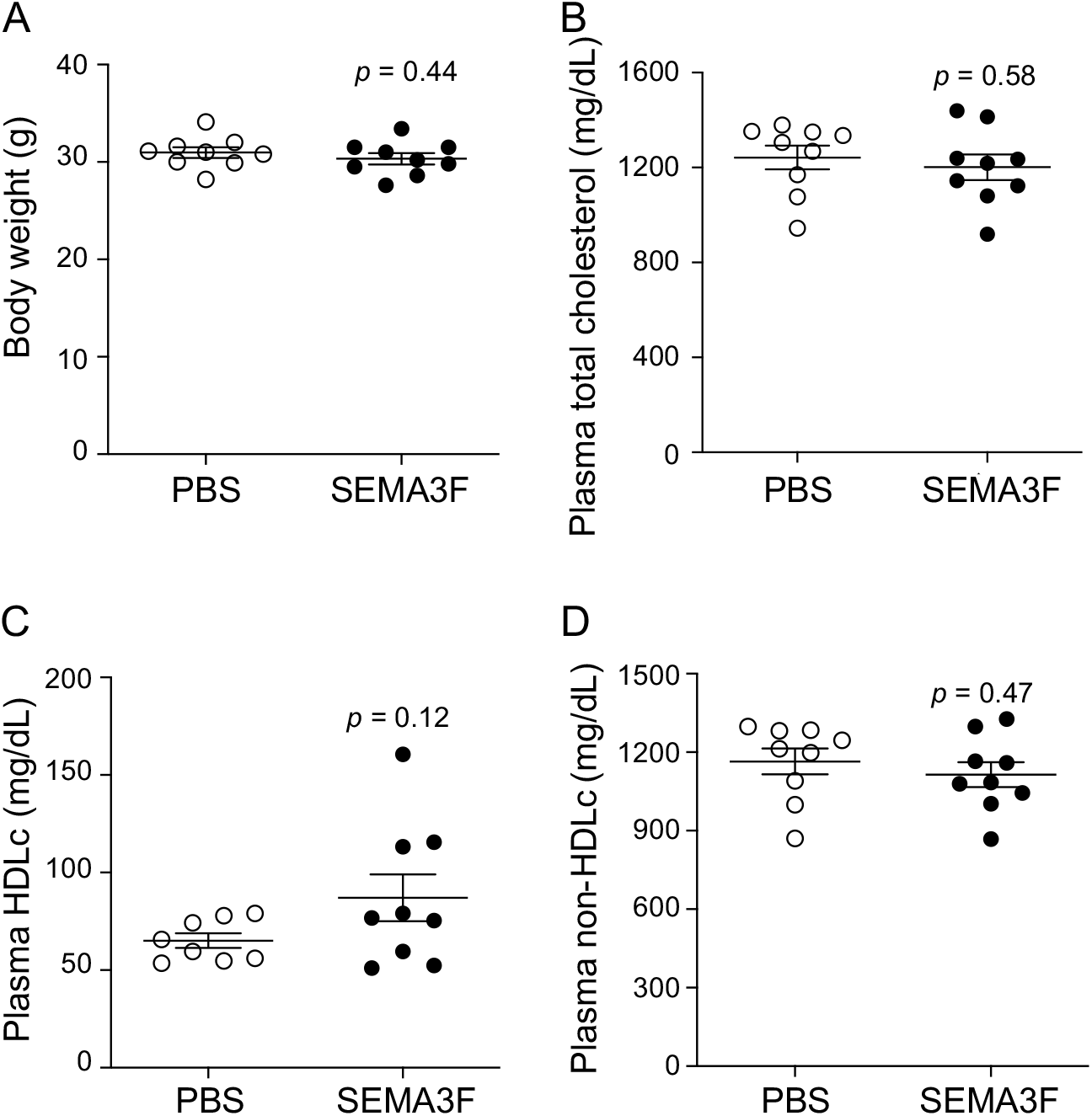
Unaltered plasma lipoprotein levels in mice administered SEMA3F. **(A)** Body weights are unchanged in western-type diet-fed *Ldlr^-/-^* mice administered SEMA3F. Plasma **(B)** total cholesterol, **(C)** HDL cholesterol, and **(D)** Non-HDL cholesterol levels were unaltered in western-type diet-fed *Ldlr^-/-^* mice administered SEMA3F. n=8-9/group. Groups are abbreviated as: PBS, *Ldlr^-/-^* mice fed western-type diet, injected with PBS; SEMA3F, *Ldlr^-/-^* mice fed western-type diet, injected with SEMA3F protein. VECs, vascular endothelial cells. Values represent mean ± SEM. All data were normally distributed and analyzed using Student’s *t*-tests.

**Supplementary Figure 5:**
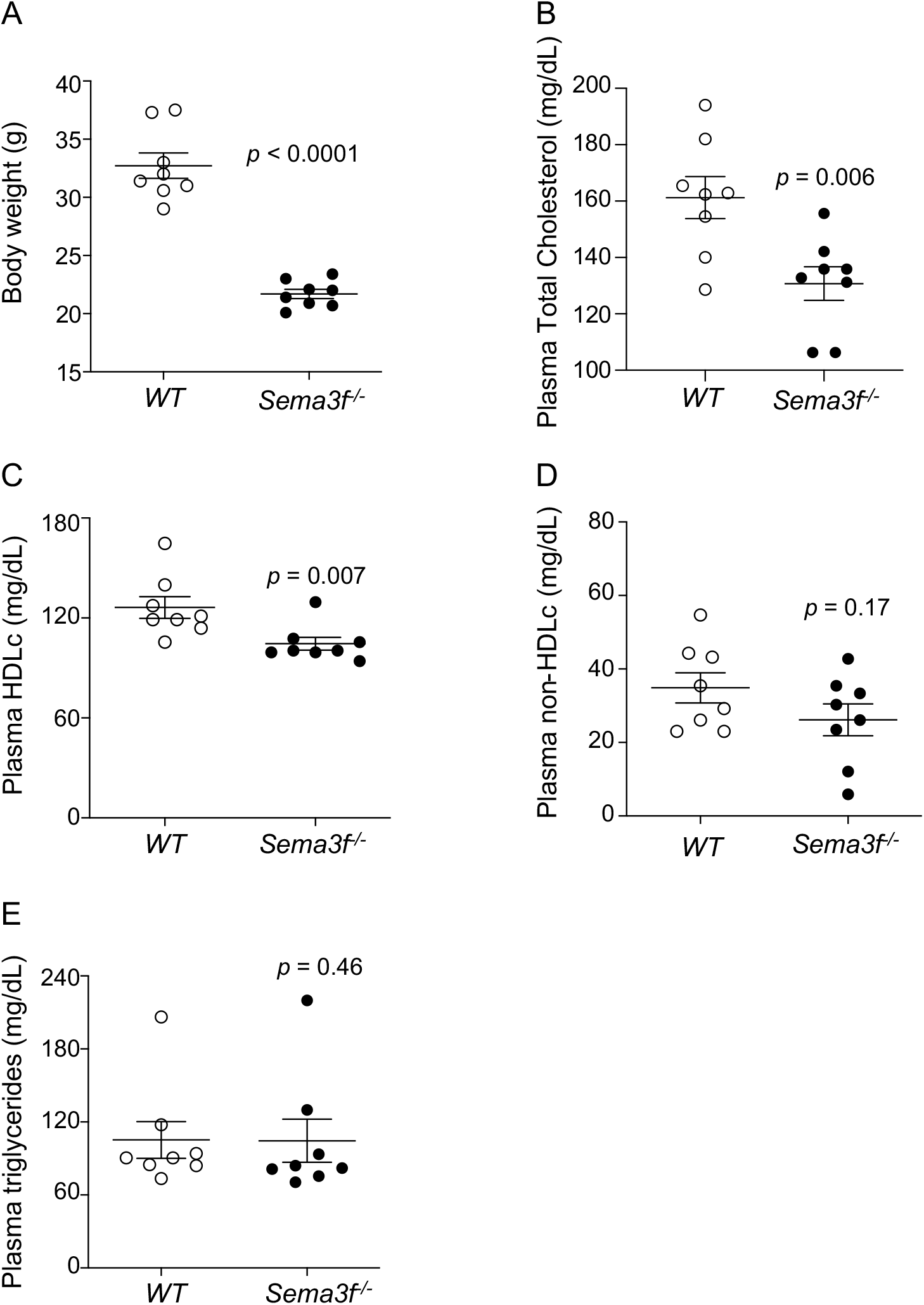
Body weights and plasma lipoproteins in *Sema3f^-/-^* mice. **(A)** Body weights were significantly decreased in *Sema3f^-/-^* mice fed a western-type diet. Plasma **(B)** total cholesterol, and **(C)** HDL cholesterol were decreased, whereas **(D)** plasma non-HDL cholesterol, and **(E)** triglyceride levels were unaltered in *Sema3f^-/-^* mice fed a western-type diet. n=8/group. Groups are abbreviated as: *WT*, wild type mice; *Sema3f^-/-^*, *Sema3f* knockout mice. Values represent mean ± SEM. Data in (A), (B) and (D) were normally distributed and analyzed using Student’s *t*-tests. Data in (C) and (E) were not normally distributed, and analyzed using Mann-Whitney *U*-tests.

**Supplementary Figure 6:**
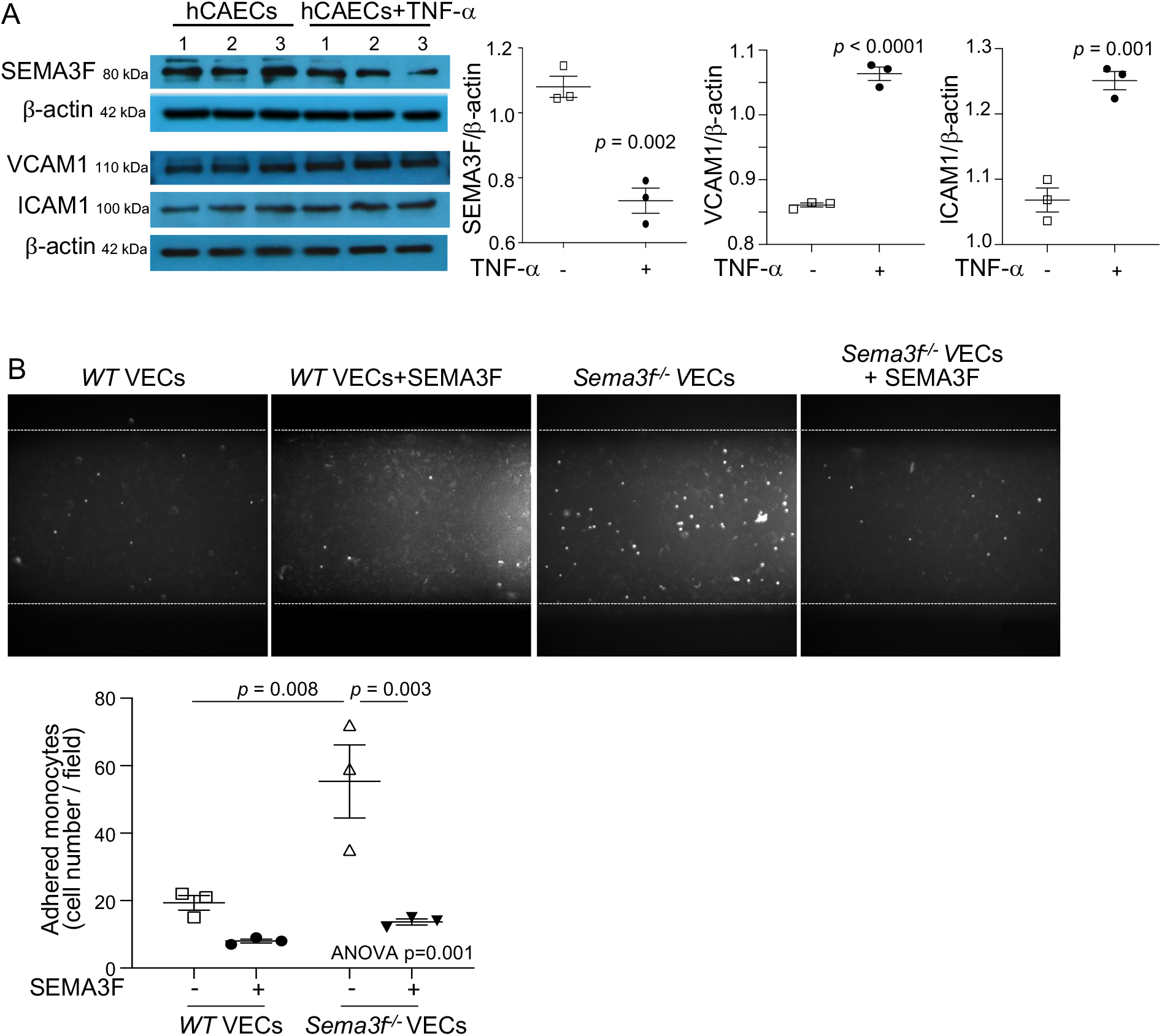
Endothelial SEMA3F directly modulates monocyte adhesion to vascular endothelial cells. **(A)** TNF-α-induced activation decreases endothelial SEMA3F, and increases VCAM1 and ICAM1 levels. **(B)** The number of human monocytes adhered to *Sema3f^-/-^* VECs was higher, and pre-treating *Sema3f^-/-^* VECs with SEMA3F protein reversed monocyte adhesion, showing that SEMA3F directly modulates endothelial monocyte adhesion. Groups are abbreviated as: *WT*, wild type; *Sema3f^-/-^*, *Sema3f* knockout mice; VECs, vascular endothelial cells. Values represent mean ± SEM, n=3/group. Data in A were normally distributed and assessed using Student’s *t*-test. Data in B was analyzed using One-way ANOVA followed by Tukey post-hoc test for multiple comparisons. Scale bar in (B): 200 µm.

**Supplementary Figure 7:**
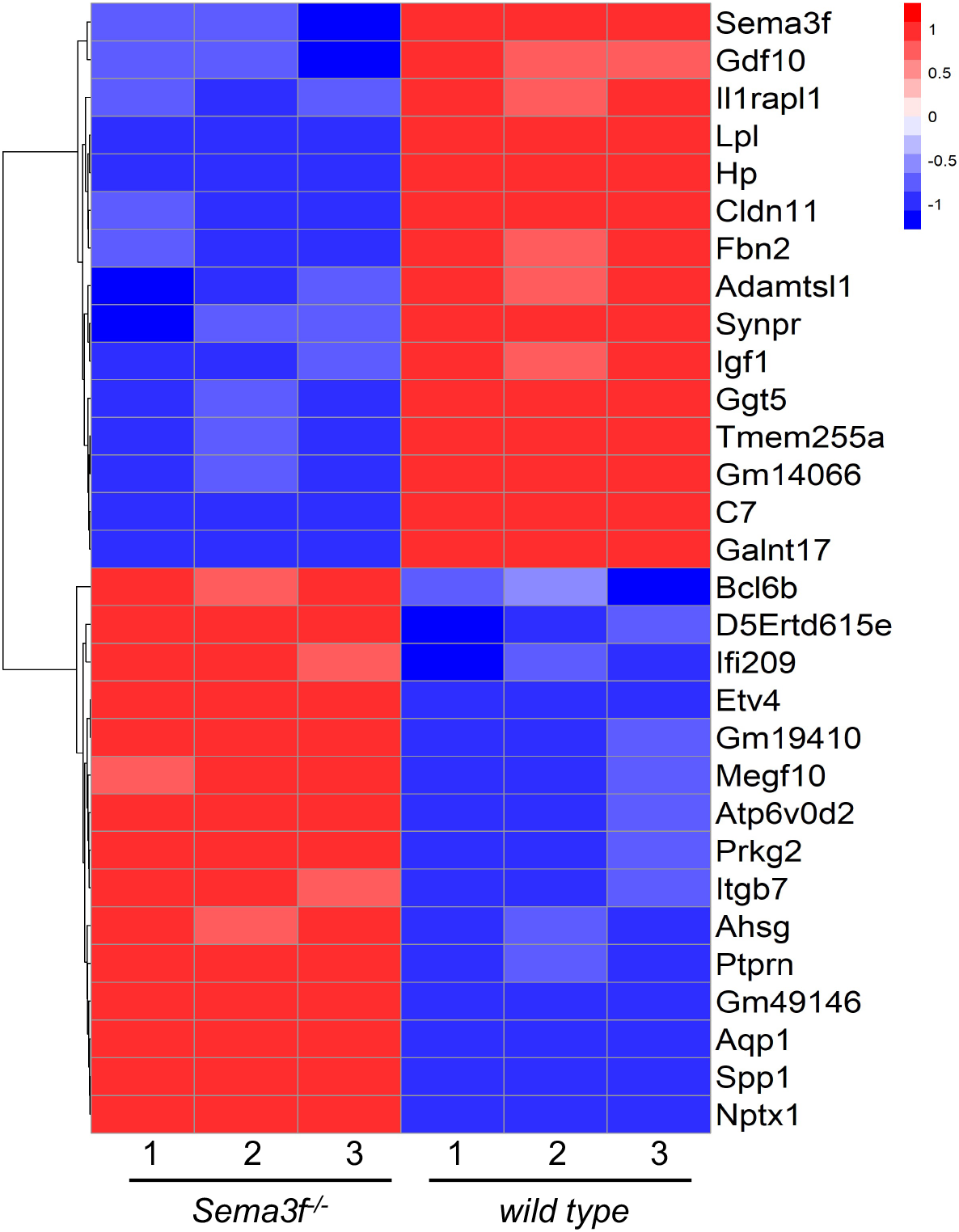
Top transcriptionally regulated genes in *Sema3f^-/-^* endothelial cells. Heatmap shows the top 30 most significantly transcriptionally regulated genes (15 upregulated and 15 downregulated) in VECs from *Sema3f^-/-^* and *WT* mice. Heatmaps are color-coded by the row-normalized log2 expression values, with shades of blues and reds representing lower and higher expression levels, respectively.

**Supplementary Figure 8:**
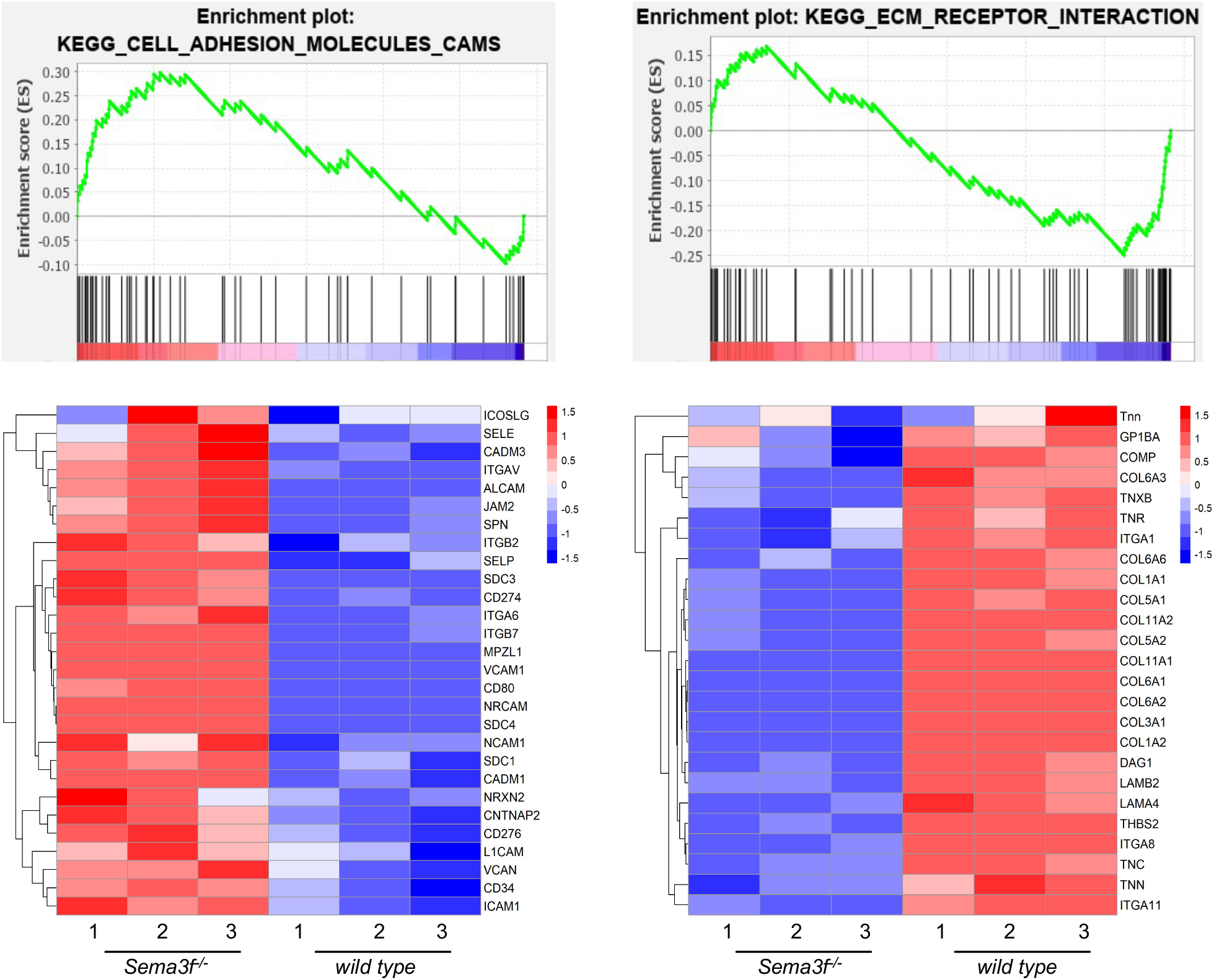
Gene-set pathway enrichment analysis in *Sema3f^-/-^* aortic endothelial cells. Gene expression data was analyzed for pathway enrichment using the KEGG pathway database via GSEA. The top panels in show the enrichment plots for the ‘*Cell adhesion molecules, CAMS’* and ‘*ECM receptor interaction*’ pathways on the left and right, respectively. These plots represent the incremental change in enrichment scores for the pathways when queried against the ranked list of genes in GSEA. The bottom panels depict the relative expression of the genes (contributing to the enrichment of these pathways) in the *Sema3f^-/-^* and *wild-type* samples respectively. All heatmaps are color-coded by the row-normalized log2 expression values, with shades of blues and reds representing lower and higher expression levels, respectively.

**Supplementary Figure 9:**
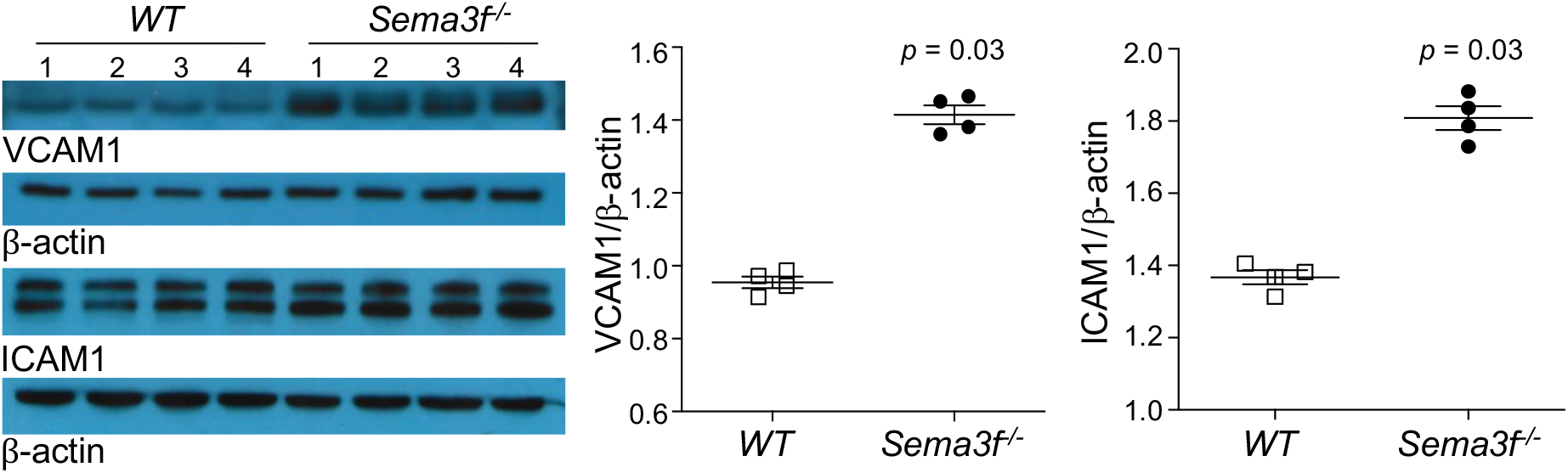
Increased VCAM1 and ICAM1 protein in *Sema3f^-/-^* aortic endothelial cells. Groups are abbreviated as: *WT*, wild type; *Sema3f^-/-^*, *Sema3f* knockout mice. Values represent mean ± SEM, n=4/group. Data were assessed using the Mann-Whitney *U*-test.

**Supplementary Figure 10:**
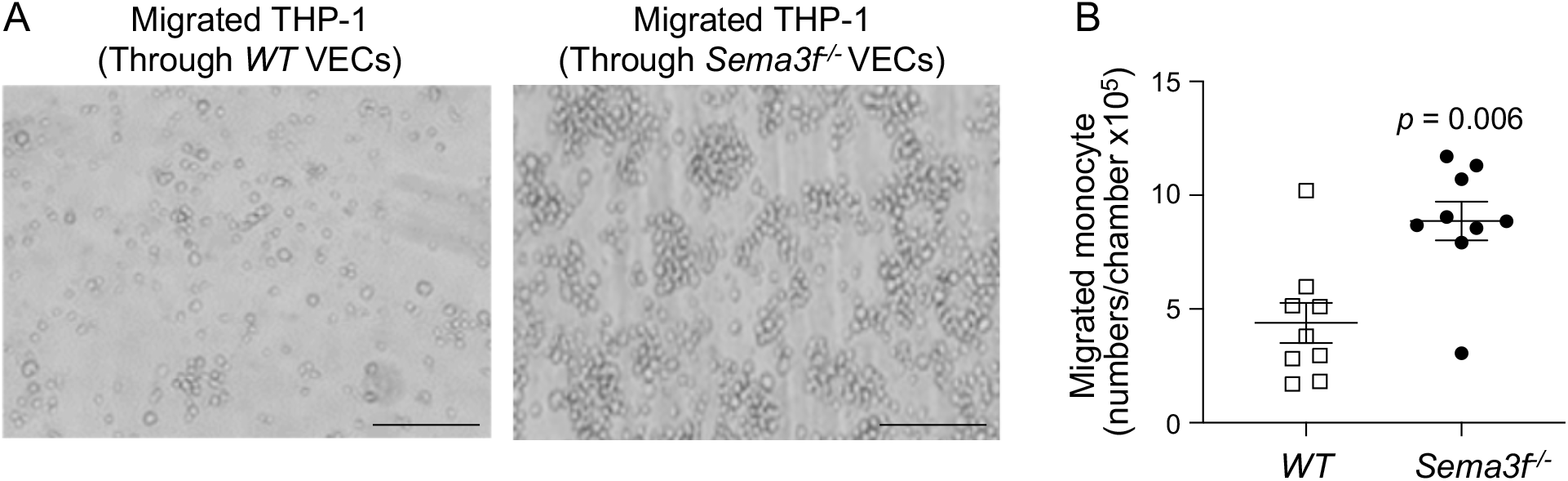
Transmigration of THP-1 monocytes through the vascular endothelial cells from *Sema3f^-/-^* mice is increased. **(A)** Representative photomicrographs of migrated THP-1 cells through *WT* and *Sema3f^-/-^* VECs. **(B)** The number of migrated THP-1 cells through *Sema3f^-/-^* VECs is higher than that through *WT* VECs. n=9 each. Groups are abbreviated as: *WT*, wild type mice; *Sema3f^-/-^*, *Sema3f* knockout mice; VECs, vascular endothelial cells. Values represent mean ± SEM. Data in (B) were analyzed using Mann Whitney *U*-test. Scale bar: 100 µm.

**Supplementary Figure 11:**
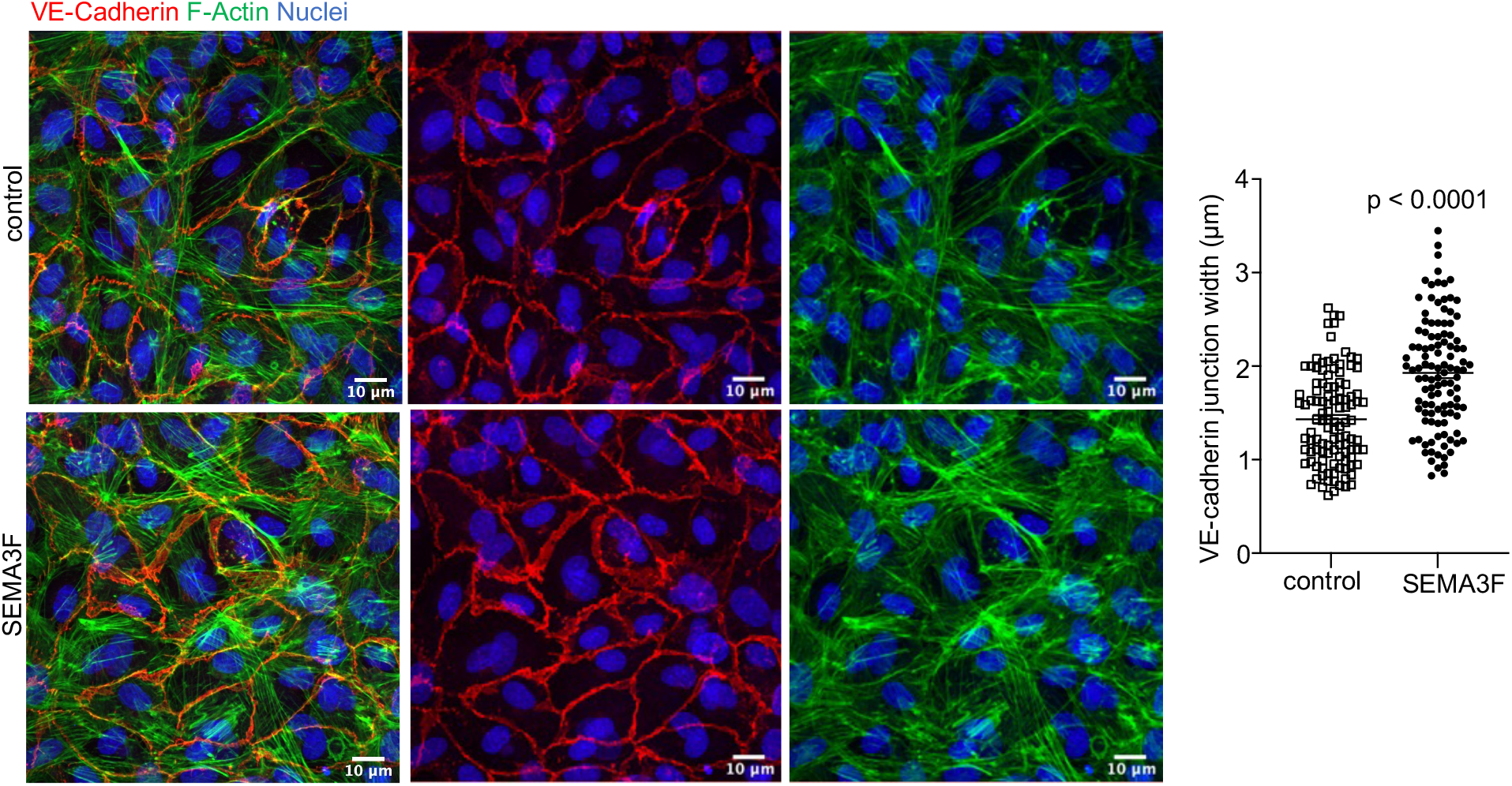
Increased adherens junction thickness in SEMA3F-treated aortic endothelial cells. Treatment of primary human aortic endothelial cells with SEMA3F resulted in significantly increased cell-cell VE-cadherin junction thickness, indicating that SEMA3F decreases vascular endothelial permeability. n=111 each. Values represent mean ± SEM. Data were analyzed using Mann Whitney *U*-test. Scale bar: 10 µm.

**Supplementary Figure 12:**
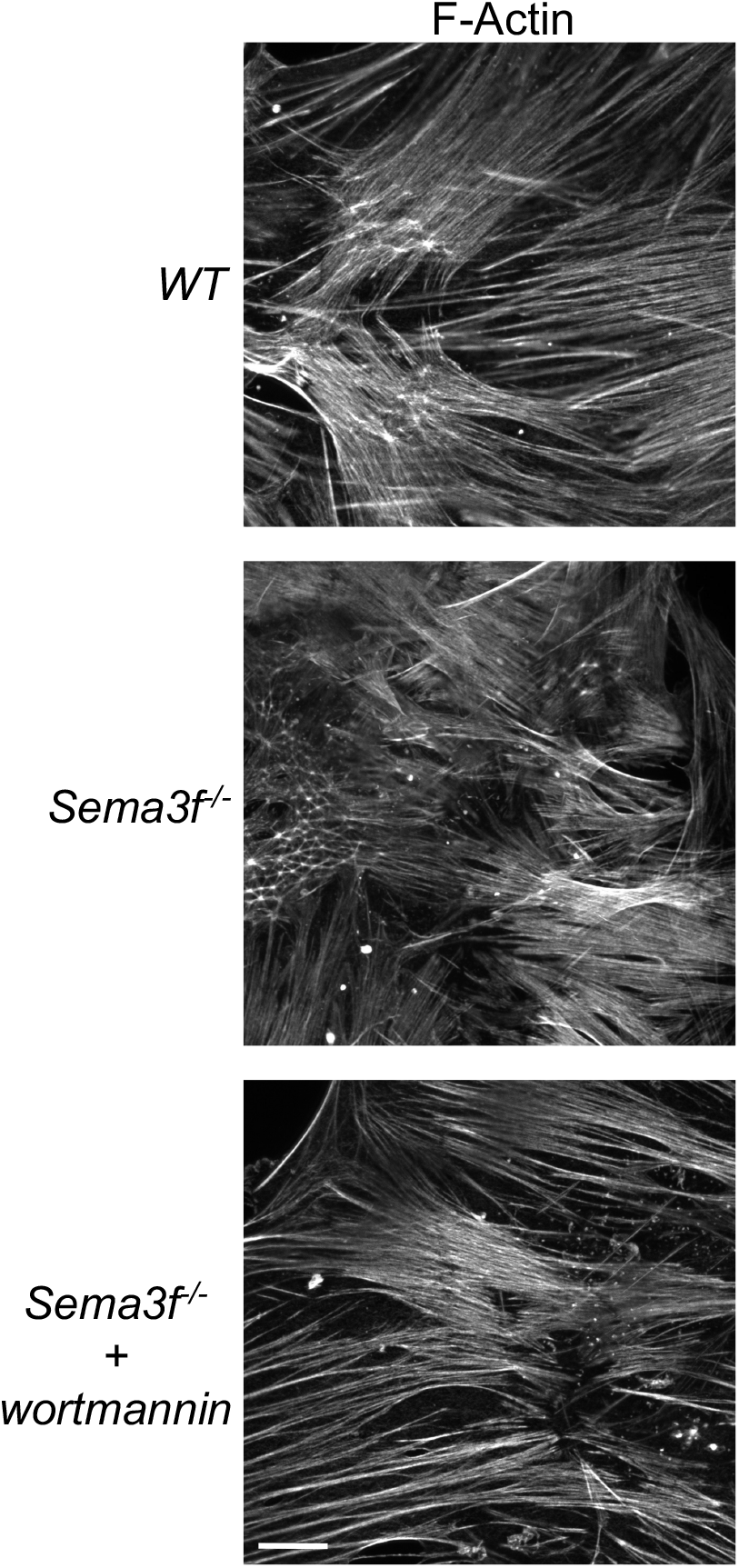
Increased PI3K in the absence of SEMA3F regulates endothelial F-actin stress fibers. F-actin fibers are decreased in aortic endothelial cells from *Sema3f^-/-^* mice, and this decrease is normalized upon treatment of the *Sema3f^-/-^* endothelial cells with the PI3K inhibitor wortmannin, suggesting that SEMA3F modulates F-Actin stress fibers via decreasing PI3K activity. Scale bar: 10 µm.

**Supplementary Figure 13:**
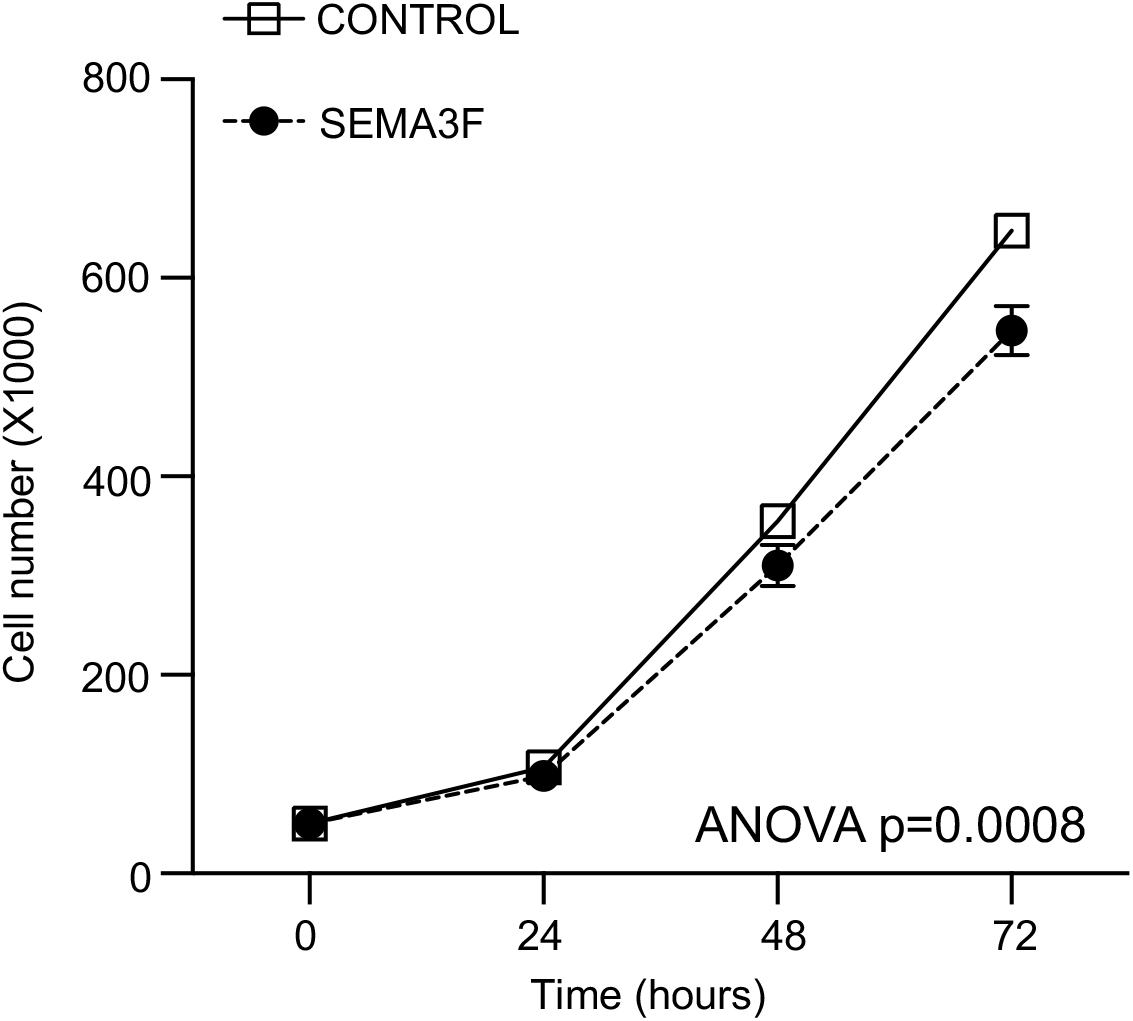
SEMA3F decreases proliferation of human coronary artery smooth muscle cells. The proliferation of HCASMCs was decreased upon exposure to SEMA3F as quantified by cell counting at various time-points after plating. n=3 each. Values represent mean ± SEM. Data were analyzed using Two-way ANOVA.

